# Asymmetric nature of MscL opening revealed by molecular dynamics simulations

**DOI:** 10.1101/2025.02.10.637407

**Authors:** Olga N. Rogacheva, Wojciech Kopec

## Abstract

We employed all-atom unbiased molecular dynamics to simulate the L17A, V21A mutant of MscL. Under a tension of 30 mN/m, the closed state adopts a funnel-like conformation. Subsequently, five chains of MscL undergo sequential transitions, leading to the formation of the asymmetric states (S1, S2, etc.). After the first chain crossed the barrier, the next transition favors the neighboring clockwise chain. This study focuses on the S1 state. Within its "open" fragment, this state is similar to the expanded state of M. acetivorans MscL, and has a conductance 10 times lower than the open state. We applied committor analysis and a nonlinear regression model to construct a reaction coordinate for the transition between the closed and the S1 state as a linear combination of interatomic distances and contacts. The main contributions to the reaction coordinate are: 1) the disruption of the "cytoplasmic" contact sites between the considered chain and two adjacent chains 2) the delipidation of the lipid-binding pocket, formed by the I82, V86, and V22 residues. The result is the pulling of the considered chain via the "cytoplasmic" tension sensor. The free energy profile along the reaction coordinate was calculated using the umbrella sampling approach. The S1 state is approximately 5 kJ/mol more favorable than the closed state under tension, and the height of the free energy barrier for the transition towards the S1 state is approximately 10 kJ/mol. This barrier is in a reasonable agreement with the corresponding average transition time, estimated to be 133 ± 13 ns.

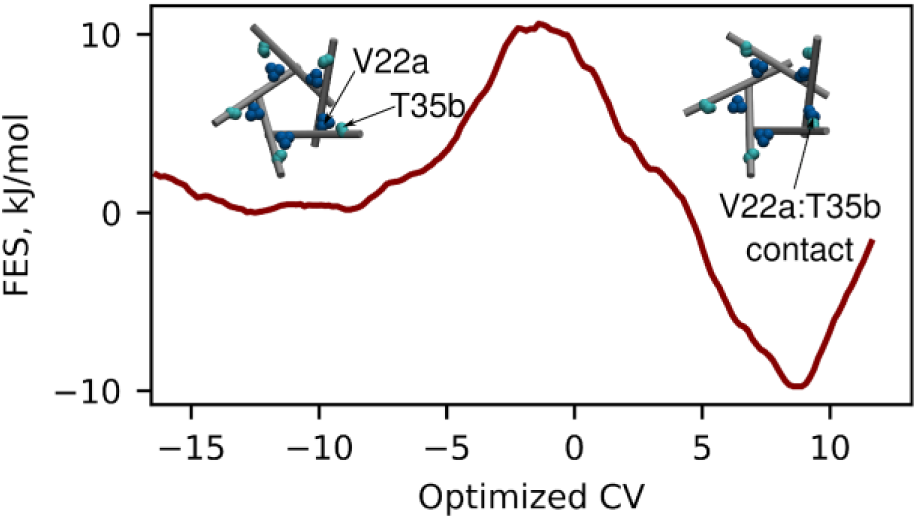

## INTRODUCTION

The mechanosensitive large-conductance channel (MscL) is one of the bacterial channels that responds to changes in membrane tension during osmotic shock. It serves as a final line of defense by opening at exceedingly high tension thresholds, approaching the membrane lytic limit, particularly at 10–12 mN/m for E. coli, and at tensions that are even twice as high in M. tuberculosis^1,2^. When open, the channel forms a large pore of 30 Å diameter^1^, which allows ions and other valuable small components to leave the cell in a non-selective manner, but effectively prevents cell rupture.

The primary stimulus responsible for the gating of the MscL is the membrane tension^3^. Nevertheless, additional factors have been shown to influence this response. For example, membranes containing lipids with shorter tails have been observed to exhibit a diminished tension activation threshold^4^. Furthermore, the presence of lysolipids in the periplasmic membrane leaflet^5, 6^, and specific mutations of membrane-facing residues of MscL, such as L89W^7^, can even result in spontaneous channel opening at zero membrane tension. These observations indicate that not only membrane tension itself, but also more subtle interactions between the protein and the lipids, play a role in MscL gating.

A crystallographic analysis of the closed state of M. tuberculosis MscL (MtMscL) has revealed that the protein is composed of five identical chains^8,9^ (Figure 1a). Each chain is composed of a short amphipathic N-terminal helix, situated in the membrane in close proximity to the intracellular side, two transmembrane helices (designated as TM1 and TM2) connected by a periplasmic loop, and a C-terminal cytoplasmic helix. The TM1 helix of one chain and the TM2 helix of the adjacent chain, which is in a clockwise position when viewed from the periplasmic side of the channel, interact with each other by multiple contacts, forming a rib of the channel that is highly stable and unlikely to disassemble during the MscL opening process^10^. In this rib the TM1 helix lines the pore, and the TM2 helix faces the membrane. A multitude of experimental and computational studies have demonstrated that the five MscL ribs undergo a tilting and shifting motion relative to one another in response to high tension^11–16^. In its closed state, the pore of the MscL exhibits a hydrophobic constriction in close proximity to the intracellular side of the membrane. This constriction is constituted by the two rings of the L17 and V21 side chains (see Figures 1a and 1b).

**Figure 1.**
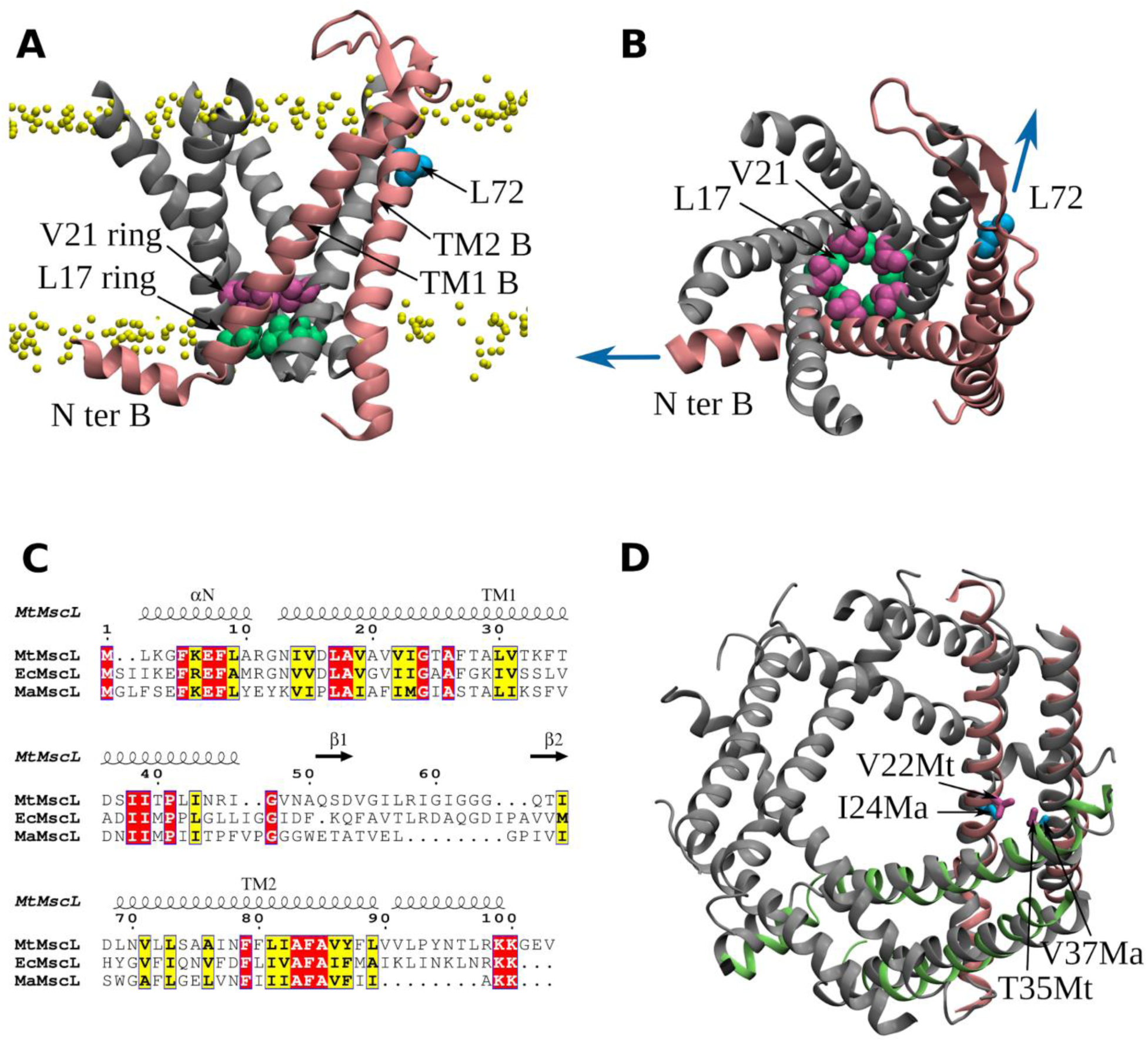
The structural characteristics of the MscL. **A.** The crystal structure of the closed state of MtMscL (PDB: 2OAR), with the cytoplasmic C-terminal domain removed. One of the five identical chains (chain B) is highlighted in pink. In the remaining chains, only the TM1 helices are shown. The border between polar lipid heads and nonpolar tails is demonstrated with yellow beads. **B.** The same closed state, viewed from the periplasmic side of the membrane. The approximate directions of tensile forces applied to the tension sensors of the chain B are illustrated with blue arrows. **C.** The protein sequence alignment of MtMscL, EcMscL, and MaMscL^19^ is depicted with the Espript 3.0 web server^20^. It is important to note that there are likely some inaccuracies in the alignment of the TM2 helix. For example, residue L89 from the MtMscL sequence, but not residue F88, should be aligned with residue M94 from the EcMscL sequence^19^. **D.** Alignment of the expanded state of MaMscL (PDB: 4Y7J) (gray) with two adjacent ribs from the MtMscL S1 state (lime and pink). The S1 state is obtained from simulations and is characterized by the displacement of the lime rib,compared to the closed state. The new position of the lime rib allows V22 and T35 residues to contact each other. I24 and V37 are homologous to these residues in MaMscL.

These rings display high resistance to wetting, thereby forming a "vapor lock"^16^. The disruption of the "vapor lock" is regarded as a rate-limiting step in the opening of the MscL pore^11^.

The process of MscL opening was elucidated with the identification of two tension sensors (Figures 1a and 1b). The initial sensor is represented by the amphipatic N-terminal helix^17^ and is situated on the intracellular side of the membrane. The second sensor in MscL obtained from E. coli (EcMscL) is formed by the residue F78, which is situated in close proximity to the outer surface of the membrane^18^. Based on the amino acid sequence alignment (Figure 1c), the homologous residue in the MtMscL sequence is L72. Given the pentameric structure of MscL, it contains a total of five periplasmic and five intracellular tension sensors. The elegant experiments involving the incorporation of lysolipids into the distinct leaflets of the membrane^6^ and the subsequent simulations that elucidate the experimental outcomes^5^, allow for the assumption that the intracellular tension sensor plays a pivotal role in MscL gating. This is presumably due to the fact that tensile forces can be effectively transferred to the hydrophobic constriction because it is located in close proximity.

Since the discovery of MscL, considerable attention has been devoted to the identification of mutants that, in one way or another, affect the functioning of the MscL. These mutants can be classified into two distinct groups. The first set of mutations is postulated to disrupt the protein structure, leading to a loss of function (LOF). Consequently, the channel containing such mutations is unable to undergo activation by tension. The second group of mutants, designated as "gain-of-function" (GOF), exhibit an inverse phenomenon, displaying a lowered tension threshold. This can be attributed to the lowered free energy barrier on the path of MscL opening. Both LOF and GOF mutants elicit an inappropriate response of MscL to the tension stimulus, which ultimately results in a decreased cell viability^21^. The use of gain-of-function mutants can provide significant insight into the understanding of MscL gating. In particular, GOF mutants can be valuable in molecular dynamics studies, as they can markedly reduce the necessary simulation time. The majority of mutations that result in the gain-of-function (GOF) phenotype are situated along one side of the TM1 helices, which lines the pore. The most severe of these mutations are those that render the hydrophobic constriction^22,23^ more susceptible to wetting. For instance, a GOF phenotype was observed in the L17Y and V21A mutants^22^, and another study determined that the V21A mutant exhibited a twofold reduction in the tension threshold^2^.

A variety of computational methods have been employed to model the process of MscL opening. The seemingly straightforward approach of applying tension to the membrane is, in fact, a challenging and often unsuccessful method. This is due to the fact that the MscL channel opens at a tension level close to the membrane lytic limit and requires a long waiting time before the opening event occurs. It is possible that the membrane may rupture before the channel opens. Nevertheless, there have been successful attempts to obtain expanded MscL structures with a large pore using this approach in both coarse-grained and full-atom simulations^11,12,24,25^. Furthermore, some of these studies employ GOF mutants and demonstrate that hydrophilic substitution of residues implicated in "vapor lock" can markedly accelerate the MscL opening process^11,12^. An alternative approach is to apply external pulling forces to the MscL atoms within the context of steered molecular dynamics simulations. In this approach, forces were applied to all atoms, to selected atoms, or to the N-terminal tension sensor^26,27,28^. In all cases, MscL underwent a transition to a structure with a large pore. Recently, an elegant approach to induce MscL opening has been proposed, whereby forces are applied not to the protein but to lipids^13^. Similarly, the conducting state with a large pore was obtained. However, the lack of structural data and a detailed description of the contacts between residues in the modeled states with a large pore makes it difficult to compare them with each other and with the available experimental data obtained for MscL in its open state. The situation is further complicated by the paucity of experimental data on the open state that can be directly converted into structural information. Among the available data are a few detected contacts between residues resulting from disulfide trapping experiments and a set of distances derived from FRET experiments^10,15,29,30^. The sole spatial structure of the non-closed state of MscL is that of Methanosarcina acetivorans (MaMscL), which represents an intermediate state rather than an open one^14^.

In the present study, we employed the L17A,V21A GOF mutant of MtMscL and conducted simulations without external forces, applying a tension of 30 mN/m, which is much lower than in previous studies (40 mN/m and above)^11,12,24,25^. This approach permitted the observation of multiple spontaneous transitions between the closed state and the subsequent intermediate state, designated as S1. The S1 state is notable for its asymmetric nature, as evidenced by the observation that only one rib assumes a position that is nearly identical to that seen in the expanded MaMscL structure^14^ (Figure 1d). The intrinsic asymmetry of the MscL intermediate states has been previously reported in experiments, in contrast to the mechanosensitive channels of small conductance (MscS), which demonstrate a completely symmetric opening. It was discussed that asymmetric intermediates must be considered in models of MscL gating^31,32^, and our study represents the first attempt to develop such a model. We conducted a comprehensive analysis of the transition between the closed and S1 states. By employing the multiple transition pathways and committor analysis, we identified a transition state region and optimized a collective variable that could distinguish between the closed and S1 states, as well as the transition state between them. Additionally, using the optimized collective variable, we estimated a free energy profile along the transition between the closed and S1 states.

## METHODS

### Molecular dynamics simulations

Wild-type M. tuberculosis MscL is known to have a very high membrane tension activation threshold^2^, exceeding 20 mN/m. On the other hand, MscL with single GOF mutations of L17 and V21 residues are more readily activated^23^. We hypothesised that the L17A,V21A double mutant might have an even lower activation tension threshold and, as implied for all GOF mutants, share a common activation mechanism with the wild-type channel. The employment of such a mutant would facilitate the simulation of MscL conformational transition, thereby elucidating the factors that are crucial for both mutant and wild-type MscL opening. With this in mind, we prepared the L17A,V21A double mutant of the MscL channel. The mutations were introduced into the MscL structure with PDB ID 2OAR. We deleted the C-terminal 5-helix bundle domain (residues 104- 125), as this domain has been shown not to be essential for MscL activation^33^. We inserted the protein into the lipid bilayer using the CHARMM-GUI server and placed the whole system in the box (15x15x12 nm) filled with 1M KCL solution^34^. The lipid bilayer contained 595 POPC lipids (300 in the lower bilayer and 295 in the upper bilayer). All simulations were conducted with the Gromacs 2020 and Gromacs 2022 software packages, using the Charmm36m force field and the TIP3Pm water model^35–37^. The NγT ensemble (constant surface tension) was employed with the Berendsen pressure coupling algorithm, which is recommended for constant tension simulations in Gromacs^38^. The temperature was maintained at 320 K through the stochastic velocity rescaling thermostat. The MscL structures were rendered using the Visual Molecular Dynamics (VMD) program^39^.

We started with 12 replicas of the MscL mutant and equilibrated them at zero surface tension for 200 ns (the workflow of a single replica simulation is illustrated in Figure 2a). Subsequently, the surface tension was increased to 10 mN/m, and the equilibration process was continued for an additional 100 ns. Finally, the systems were equilibrated at 30 mN/m surface tension for a further 50 ns. From each of these equilibrated systems, ten production trajectories, each 400 ns in length, were initiated at 30 mN/m surface tension, with the objective of observing spontaneous channel opening. However, in contrast to our initial expectations, we observed that only two adjacent ribs of the channel changed their position relative to each other. This transition occurred within a timespan of no more than 300 ns, and the resulting state (S1) did not undergo any further transitions for approximately 100 ns on average. The S1 state was obtained on at least one occasion for each set of ten trajectories. One of the S1 states was selected for each set and the next ten trajectories, each 400 ns in length, were recorded from each selected state. The subsequent conformational changes occurred in at least one instance among each set of ten trajectories, resulting in the formation of a new state (S2), which, like S1, was stable for a short period of time. This novel transition involved two adjacent channel ribs (at least one of which was distinct from those involved in the first transition) and occurred in approximately 200 nanoseconds.

**Figure 2.**
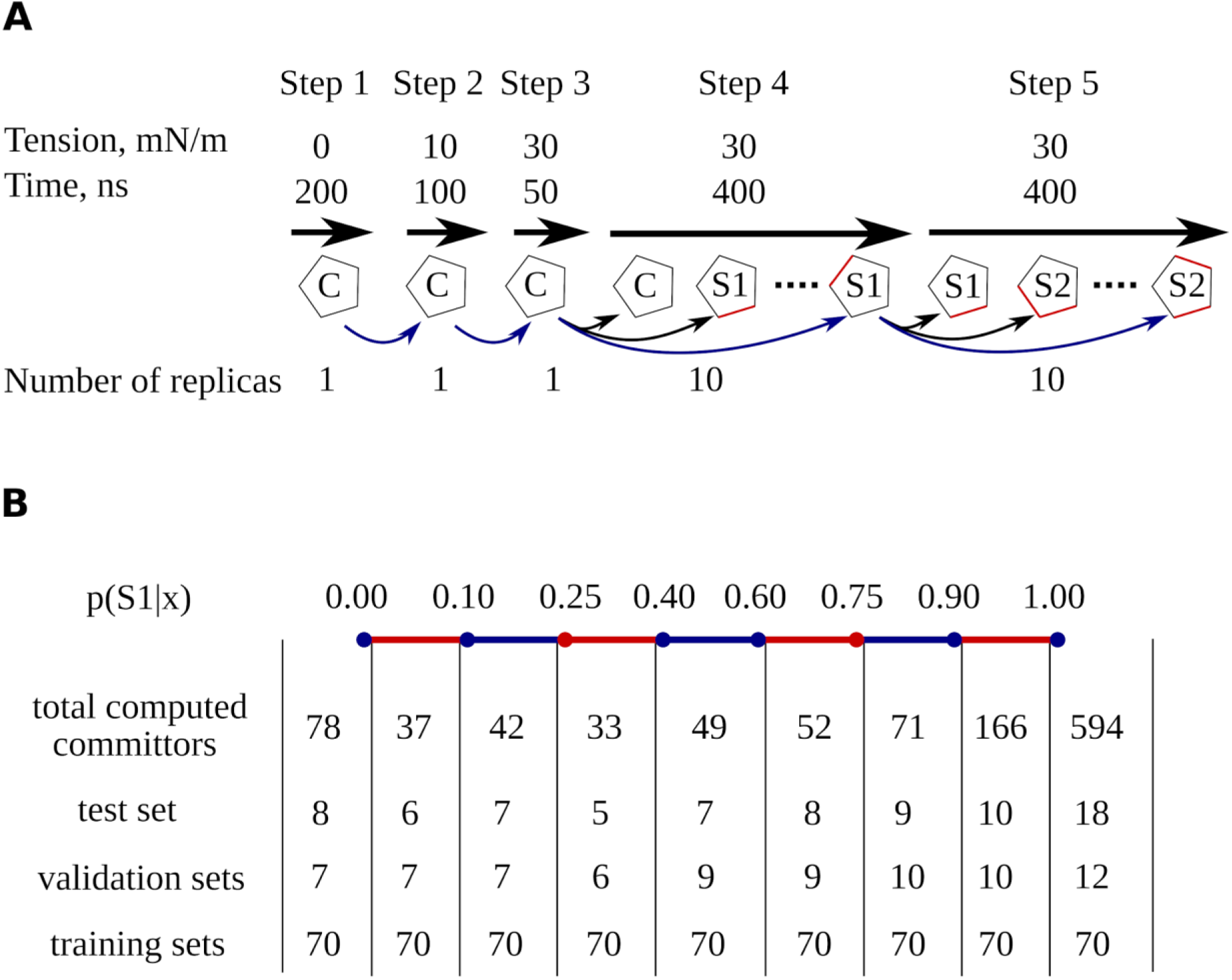
Explanations of modeling and analysis of MscL gating. **A.** the workflow of a single replica simulation. Each pentagon represents a single replica of MscL in one of three states: closed (C), S1, or S2 state. The ribs affected by transition are highlighted with red. All simulated paths are shown with curved arrows, but for the only path chosen for the analysis the arrows are colored with blue. **B.** the distribution of committor values across the entire set of snapshots for which committors were computed, as well as for the test, validation and training sets. The large number of committor values in the training sets is attributed to the application of the bootstrap resampling technique.

The preliminary concept was to generate a single sequence of transitions for each of the 12 original replicas (as designated in Figure 2a), thereby ensuring that the transitions were entirely uncorrelated. However, there are two exceptions to this. Firstly, the two paths of Replica 7 from the closed state to S1 exhibited such marked differences that we elected to retain both. Secondly, for Replica 12, the initial and subsequent transitions occurred in rapid succession during the first production run (step 4 in Figure 2a). As a result, both paths were processed. Consequently, the initiation of 12 replicas in a closed state resulted in the identification of 14 distinct pathways leading to the S1 state. In the final stage of the simulation (step 5 in Figure 2a), the number of replicas was reduced to the original 12.

### Features selection

In order to provide a more comprehensive characterization of the simulated transitions from a structural and thermodynamic perspective, it was necessary to identify a reaction coordinate (collective variable) that connects the closed and S1 states and goes through the transition state between them. The most straightforward approach to constructing such a variable is to identify the full set of key features (such as distances and contacts between residues) influenced by the transition and to determine the linear combination of these features that best satisfies the aforementioned requirements.

The initial feature selection was based on the assumption that there should be at least a few contacts between MscL residues that are typical of the closed state at zero tension, remain stable as the protein attempts to adapt to the decreasing membrane thickness under tension, and break just as the protein is on the transition path between the closed and S1 state. In order to identify these contacts, the trajectories recorded at zero tension were processed and all contacts between residues were calculated. Two residues were considered to be in contact if the minimum distance between any two heavy atoms of the respective residues was less than 4.5 Å. Given that MscL exhibits a fivefold symmetry in its closed state, we did not differentiate between the various chains. For instance, if a contact is observed between pairs of chains (e.g., A and B, B and C, C and D, D and E, and E and A) in a single frame, that frame was counted as representing five distinct instances of that contact. Residue pairs that were in contact in at least 70% of all possible instances (number of frames multiplied by five) were subjected to further analysis.

The objective was to ascertain whether the extracted contacts could provide an initial approximation of the reaction coordinate between the closed and S1 states. To assess this, we calculated the time series of the contacts for all the trajectories connecting the closed and S1 states under the applied tension (as detailed in section 2.1, this was done for 14 such trajectories) and performed principal component analysis (PCA) on the resulting time series. A visual inspection of all the trajectories revealed that the projections of the trajectories onto the first principal component exhibited a pronounced change almost always when adjacent ribs of MscL moved apart (Figure S1 a,e). This observation provided compelling evidence that the first principal component (PC1) can approximate the conformational transition from the closed to the S1 state.

The subsequent step was to expand the set of contacts to encompass all relevant contacts pertaining to the conformational transition from the closed state to S1. To this end, we computed time series for all contacts between protein residues, as well as between surface protein residues and lipids for all trajectories. Utilizing PC1 as a template for the transitions, we then selected only those contacts that exhibited a correlation with PC1, as indicated by an absolute value of the Pearson correlation coefficient exceeding 0.25.

In order to conduct a more detailed examination of the selected contacts, we have employed an alternative definition of "contacting residues". This definition is proposed in the community-developed open-source library PLUMED^40^ and appears there under the name "coordination". It is utilized throughout the remainder of this paper:

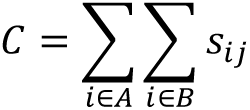

where C represents a coordination number between contacting residues A and B. The i and j variables denote heavy atoms belonging to these residues, while s_ij_ is a rational switching function that ensures the coordination number has continuous derivatives. The following formula was used for s_ij_:

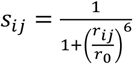

where r_ij_ represents the distance between atoms i and j, with r_0_ set to 0.3 Å.

Given that MscL opening is associated with significant alterations in pore diameter, we postulated that it could not be adequately described by short-range contacts between adjacent residues alone. To account for longer-range changes, we initially decided to calculate the distances between all CA atoms. However, to reduce the number of features without losing important information, we took advantage of the fact that MscL is a nearly helical protein and used the centers of mass of all helical turns rather than the CA atoms. We filtered the distances using the same criteria (the absolute value of the Pearson correlation coefficient between the feature and the previously defined PC1 should be greater than 0.25). All the features, such as contacts, distances, and coordination numbers, were calculated using the PLUMED library, version 2.8^41^.

### Building reaction coordinates

In accordance with the two sets of features described in section 2.2, two reaction coordinates were constructed: one comprising all relevant contacts, and the other containing all distances. Both reaction coordinates are linear combinations of the scaled features. In order to determine the coefficients associated with each feature, a methodology was employed comprising the following steps: firstly, principal component analysis (PCA) was applied to time series data obtained for all filtered features across all paths connecting the closed and S1 states; secondly, the paths were projected onto the first few principal components; and thirdly, support vector machine (SVM) classification was subsequently applied to the determined first few PCs. The analysis was conducted on the contact and distance data sets separately. When PCA was applied to the contacts, the first two principal components were retained, with a total variance of approximately 0.4. When the distances were analyzed, five principal components were retained, with a total variance of 0.75. The inclusion of additional principal components in some instances led to erroneous signs of feature weights in the final SVM-based collective variables (e.g., the contact disruptions during the transition with a positive sign), corroborating the notion that protein atoms engage in diverse movements and not all of them are associated with the transition from a closed state to S1.

As the SVM classifier is a supervised learning method, it was necessary to make an initial determination as to which frames belong to the closed or S1 states. In order to achieve this, we projected all paths onto the PC1, which is described in Section 2.2, and employed a two-component Gaussian mixture model for the differentiation between the closed and S1 states. As the PC1 did not provide a unified transition state region, the Gaussian mixture model was applied to each path separately. The separation was deemed successful if the data did not overlap within a two-sigma interval. However, this criterion was not met for all paths (e. g. the path colored in gray in Figure S1 a, e), necessitating the adoption of an alternative approach. Considered this evidence, we applied PCA to the time series data obtained for all filtered contacts. The first principal component of the all-contacts PCA was capable of differentiating between the closed and S1 states across all paths (Figure S1 b, f). Accordingly, the all-contacts PC1 was utilized for the classification of the path frames between the closed and S1 states. The classification approach employed was identical to that previously described in this paragraph, including the application of a Gaussian mixture model to each path separately. This was done because the new PC1 also failed to identify a unified transition state region.

The SVM algorithm was applied separately to distances and contacts, yielding two reaction coordinates. However, a single reaction coordinate was required for reasons related to the evaluation of the quality of the transition state. With this in mind, the optimal linear combination of the two SVM-based variables was constructed using committor values, as detailed in Section 2.4. The projections of the paths onto the combined SVM-based collective variable are illustrated in Figures S1c and g. As illustrated in Figure S1 g, the graph of the configuration densities along the collective variable reveals that the low-density region falls within a narrow range of collective variable values for all paths. It can thus be postulated that this region may correspond to the putative transition state, and that the SVM-based collective variable is capable of differentiating between closed and S1 states. The only deviation from the transition state position common to the majority of the paths was demonstrated by Replica 4 (Figure S1 g, red path). Upon visual examination of this trajectory, it was observed that flexible loops connecting the TM1 and TM2 helices were stuck together at the mouth of the channel (Figure S1 g, inset). Throughout the trajectory, they attempted to disentangle themselves and eventually succeeded. As it was uncertain whether this event would significantly alter the free energy landscape of the MscL opening, this pathway was excluded from further analysis.

### Optimization of the reaction coordinate

As has been previously stated, the SVM-based distances and contacts collective variables are nearly capable of differentiating between the closed and S1 states. Consequently, they provide the opportunity to conduct a committor analysis and further optimize the reaction coordinate. It is expected that the optimized coordinate will demonstrate the capacity to distinguish not only between the aforementioned states, but also to indicate a region of the transition state.

A committor analysis was conducted using the Plumed 2.8 library^40,41^. The committor function is defined as the probability, p(S1|x), that a trajectory initiated at a configuration space point x will reach the S1 state before reaching the closed state. The probability for a specified protein configuration was calculated based on a set of 100 short trajectories initiated from this configuration with random velocities obeying Maxwell Boltzmann statistics at 320 K.

In order to compute committor values for all snapshots located the vicinity of the transition state and to consider each trajectory initiated from these snapshots as committed if it crosses the boundary of either the closed or S1 state, it was necessary to approximately define the borders of the closed and S1 states, as well as the reactive path regions between them. To this end, we calculated the density of configurations derived from simulated paths in the two-dimensional space defined by the SVM-based collective variables. Subsequently, the plot was examined visually to identify regions of high density, which approximately correspond to the closed and S1 states, as well as a rectangular zone of low density between them. This latter zone represents a reactive path region (Figure S2a). The values of the committor were calculated for all frames falling within the specified region, with two exceptions. First, as previously discussed, the frames from Replica 4 were excluded from the analysis. Secondly, the committor values calculated on path 11 were not reproducible with the optimized version of the collective variable. Consequently, these values were also excluded, and the optimization process was repeated once more without these values. A total of 1,122 committor values were calculated for the remaining 12 paths, and the distribution of these values is illustrated in Figure 2b. A committor value of 0 indicates that all 100 trajectories initiated from the specified configuration have been committed to the closed state. Conversely, a committor value of 1 signifies that all trajectories have reached the S1 state.

The complete process of collective variable optimization is illustrated in Figure S3. The methodology is based on two assumptions. The first assumption is that the committor function can be sufficiently approximated by a logistic function if the independent variable is an adequate reaction coordinate. The second is that an adequate reaction coordinate can be presented by a linear combination of features dependent on the system coordinates in a configuration space (in this case, distances and contacts). In view of these two assumptions, a least-squares fit of a logistic function to committor values can be performed, thereby enabling an optimal linear combination of selected features to be identified. This linear combination of features may prove more effective in distinguishing between protein states than principal components or support vector machine-based coordinates, as it is based on information pertaining not only to metastable states but also to transition paths. However, if the selected set of features lacks important factors, the effectiveness of this approach may be limited. While the optimized collective variable will still be an optimal combination of given features, it will not be the optimal reaction coordinate.

The initial set of contacts and distances exhibits a high degree of multicollinearity, which must be removed to ensure the reliability of the subsequent analysis (Figure S3 a). To achieve this, the features were first filtered, and only those that correlate with committor values with an absolute value of Pearson correlation coefficients higher than 0.1 were retained. Next, the retained features were clustered using a combination of linear correlation and the Leiden community detection algorithm as implemented in the MoSAIC package^42^. The resolution parameter gamma was set to 0.6, which yielded a total of 121 clusters. A principal component analysis was conducted on each of the clusters, and only the first principal component was retained for further analysis. As the features within each cluster exhibited a high degree of correlation, the first principal component consistently accounted for more than 70% of the total variance. The clustering was repeated on 121 new features (first principal components) and a resolution parameter set equal to 0.7. Only a limited number of features demonstrated a correlation with an absolute value of Pearson correlation coefficients exceeding 0.7. Subsequently, these features were merged using the same PCA-based procedure, resulting in a final set of 115 features. It is noteworthy that none of the features in this final set exhibited a correlation with an absolute value of Pearson correlation coefficients exceeding 0.7.

In order to fit the committor values with a logistic function, the least-squares function of the scipy.optimize module was employed^43^. However, the lack of clarity regarding the optimal set of features required to achieve the best fit renders this procedure somewhat complex. In light of these considerations, the total fitting algorithm is divided into two steps. The first step is to determine the optimal combination of features, and the second step is to optimize the coefficients of these features. To this end, the committor values were divided into two sets: a 78-points test set and a remainder set comprising 1,044 points (Figure S3 a). The test set was utilized exclusively in the second step of fitting, while the remainder set was employed in both steps. It is noteworthy that all committor sets used for training, validation, and testing the model were selected in a manner that ensured that distinct committor values exhibited near-identical probabilities within a set (Figure 2b).

The optimal combination of features was determined through the following process (Figure S3 b). The remaining 1044-point set of the committor values was divided into a validation set and a total training set. Subsequently, the total training set was resampled using a bootstrap approach on twenty occasions, resulting in the generation of twenty training sets with nearly equal probability of distinct committor values. The initial stage of the process entailed the random selection of features. Subsequently, a Monte Carlo procedure was implemented, whereby the addition or removal of a random feature was accepted or declined based on the value of the adjusted R². For each combination of features, the model was fitted twenty times on different training sets, and the adjusted R² value was estimated twenty times on the validation set. The mean value of the adjusted R² was then compared with the adjusted R² value of the previously accepted combination of features. If a new combination of features was accepted, the coefficients were examined to ascertain whether they significantly differed from zero (P ≤ 0.05). The features with P values greater than 0.05 were removed. The Monte Carlo procedure was repeated 1,000 times, and the total process of searching for the optimal set of features was repeated 100 times, resulting in 100 combinations of features with a mean length of 17 features. Ultimately, only those features that were presented in a minimum of 17% of combinations were selected for further analysis, resulting in a total of 31 features.

The coefficients of this final combination of 31 features were optimized via a similar procedure (Figure S3 c). The 1044-point set of the committor values was resampled using a bootstrap approach on 300 occasions, resulting in the generation of 300 training sets. The model was fitted 300 times on distinct training sets, and the resulting coefficient values were averaged. The R², MAE, and MSE values were estimated 300 times on the test set, and the corresponding mean values of the metrics were subsequently calculated. The projections of all paths onto the optimized collective variable are illustrated in Figures S1d and h.

The same procedure, which employs committor values to optimize coefficients for a fixed set of features, was utilized to fit a position of the single SVM-based collective variable. In this particular case, two coefficients and a free member were fitted.

All calculations, including those pertaining to free energy estimation, were performed with the utilization of an optimized collective variable. However, for the purposes of visualization and better understanding of the gating mechanism, we divided the optimized collective variable into two distinct parts. Initially, the 31 features were clustered as described above. The resolution parameter gamma was set to 0.2, resulting in a total of six clusters. Subsequently, the clusters were divided manually into two groups, and the features belonging to each group were assembled in a linear variable with coefficients determined previously when optimizing the total variable. The Pearson correlation coefficient between the two variables was found to be 0.1.

### Free energy calculations

An umbrella sampling approach was employed to estimate the free energy profile along the optimized collective variable^44^. As unbiased paths were available connecting the closed and S1 states, there was no necessity to pull the system along the collective variable. Instead, the snapshots with appropriate values of the collective variable were selected from the available paths. The methodology employed was that of a single path yielding a single free energy profile, with the final free energy profile being the average of several free energy profiles. In this specific instance, the averaging process was conducted using four distinct paths, specifically Replicas 1, 7a, 8, and 9. The selection of these paths was based on the premise that they exhibited sufficient duration within the reactive path region, thereby enabling the retrieval of snapshots from the transition state. The Plumed 2.8 library was utilized for the execution and analyzing of the biased simulations^40,41^. The simulation parameters encompass a window width of 0.5, a biasing umbrella potential magnitude of 50 kJ/mol, an equilibration time of 25 ns for each window, and a time for free energy profile calculation of 2.5 ns.

### Estimation of the average transition time

The "true" average transition time between the closed and S1 state was calculated in accordance with the following methodology^45^. For each replica, the time elapsed between the entrance into the basin corresponding to the closed state under tension and the crossing of the barrier was estimated. The statistics of the transition events should follow a Poisson distribution, and thus the time elapsed before the transition occurs is exponentially distributed. The empirical cumulative distribution function of the transition times was calculated, thereby allowing the best-fit mean transition time (τ) to be determined. In order to ascertain whether τ can be considered a "true" average transition time, it is necessary to verify that the distribution of transition times is indeed exponential. To this end, a theoretical cumulative distribution function was calculated by sampling a large set of numbers from the exponential distribution with a mean value equal to τ. Subsequently, the two distributions were compared using a two-sample Kolmogorov–Smirnov test. A high p-value indicates that the empirical distribution is exponential, and τ can be considered a "true" average transition time.

In addition to the "true" average transition time, it is possible to determine the average transition time that would be observed if the estimate of the free energy barrier were accurate. This can be done using an analogue of the Eyring-Polanyi equation:

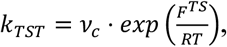

where k_TST_ represents the reaction rate constant in accordance with transition state theory, F^TS^ denotes the computed barrier height, and ν_c_ signifies the frequency with which the system attempts to traverse the barrier. The average transition time is thus equal to 1/k_TST_.

In practice, v_c_ value was estimated as the crossing frequency of 1RT level from the closed state towards the barrier. The resulting frequency was determined to be 0.85 ns⁻¹. It is important to note that this value may be an underestimate due to the infrequent storage of the trajectory, with a step of 40 ps. However, given that the transmission factor is not accounted for, the resulting average transition time can be considered to fall within the margin of error associated with such estimations.

## RESULTS

In the closed state, the five MscL ribs are in a compact arrangement and oriented almost orthogonally to the plane of the membrane (Figure 1a). In contrast, the open state is predicted to result in all five ribs lying in the plane of the membrane and moving away from each other, forming a large open pore. An approximation of the open state can be derived from the structure of MaMscL (Figure 1c). However, this state cannot be regarded as fully open, as it does not align with all experimental data (for instance, results of disulfide trapping experiments^10,30^). Instead, it can be classified as an intermediate state. The use of the GOF L17A,V21A mutant and the absence of external bias in our simulations enabled a comprehensive investigation of the opening mechanism, resulting in the identification of metastable states along the transition path. The primary observation is that, in the initial approximation, all five ribs display a notable degree of independence with regard to their movement. This results in an asymmetric opening of the channel when two adjacent ribs move apart from each other, while four other pairs of adjacent ribs can move or remain stationary. The initial intermediate state, which is defined by the separation of a mere two ribs, was designated as the S1 state. This state displayed metastable characteristics, with a transition to the S2 state occurring within a few hundred nanoseconds. This transition involved the separation of an additional pair of adjacent ribs. Subsequently, transitions into the states S3, S4, and S5 were observed. The present paper, however, aims to examine the initial transition from the closed state to the S1 state in greater detail.

For the sake of simplicity, the following notations are employed in the paper, as all five chains of MscL are initially identical. The transition from the closed to the S1 state is characterized by the separation of two ribs: one formed by the TM1 helix of chain A and the TM2 helix of chain B, and the other formed by the TM1 helix of chain B and the TM2 helix of chain C. This notation does not result in the loss of information, as the chains can always be renumerated. In the paper, this transition is also referred to as the transition of chain B, as the two helices of chain B move away from each other. Consequently, it is chain B that undergoes the most significant conformational changes when the system achieves the state S1.

### The committor-based collective variable exhibits characteristics of an optimal reaction coordinate

It was hypothesized that the optimal reaction coordinate connecting the closed and S1 states could be described by a linear combination of specific contacts between channel residues alone, channel residues with surrounding lipids, and distances between the centers of masses of specific turns of channel helices. In order to optimize the linear coefficients, we implemented a procedure that involved fitting the committor values with a logistic model (Figure S3). The reaction coordinate, when optimized in this manner, possesses the property that the transition state - which is defined by a committor value of 0.5 - is invariably projected onto the value of 0.

In order to validate the optimized collective variable, we first made sure that it can predict committor values over the entire range (Figure 3a). Further we checked the ability of the optimized coordinate to identify a transition state. A good reaction coordinate at a transition state would yield a sharp unimodal distribution of a committor values, with a mean value of p(S1|x) = 0.5. Initially, we sought to ascertain whether this was the case for the SVM-based collective variable (see Section 2.3). We took all the committor values within the interval [-0.25, 0.25] of the SVM-based collective variable and subsequently plotted the distribution of committor values within this set. In contrast to the ideal distribution, the data set exhibits a pronounced peak at committor values of 1 (Figure S2b), indicating that the SVM-based collective variable cannot identify the transition state. A similar analysis for the committor-based collective variable reveals a distinct peak in the range of 0.5 (Figure 3 b), which can be deemed satisfactory if, for reaction coordinate values greater than 0, the committor values are predominantly greater than 0.5, and for reaction coordinate values smaller than 0, they are predominantly smaller than 0.5. As demonstrated in Figure S4, the conditions under examination are indeed met, thereby substantiating the efficacy of the optimized reaction coordinate in identifying the transition state.

**Figure 3.**
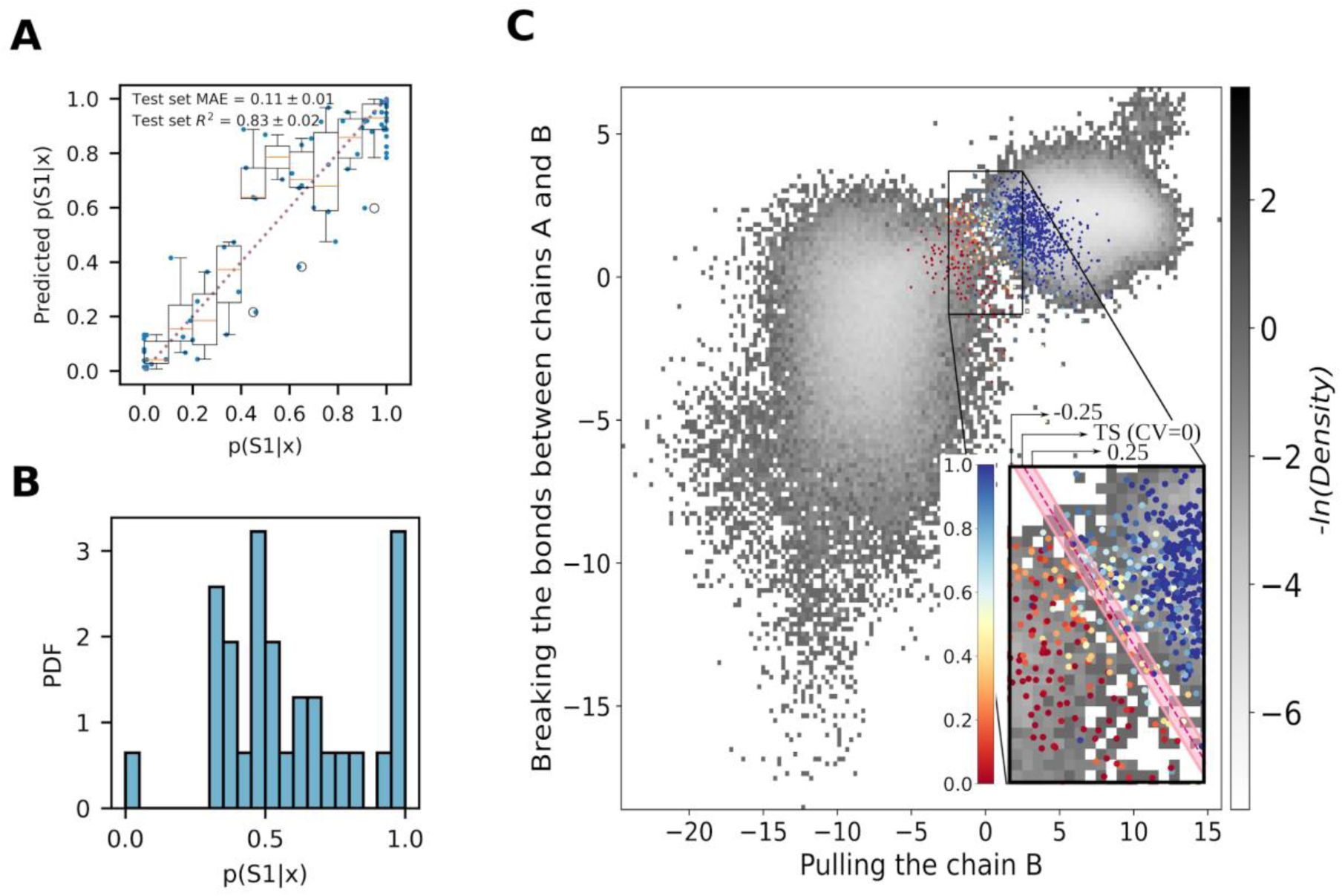
Characteristics of the optimized collective variable. **A.** A comparison of the test set committor value to the model prediction. The blue dots represent individual data points, and the box plot depicts the data partitioned into 0.1-wide bins. The orange lines represent the medians, the boxes extend from the first to the third quartile of the distribution, and the whiskers extend to 1.5 times the interquartile range. Outliers are indicated with black circles. **B.** The distribution of committor values in the transition state. **C.** The two-dimensional space derived from splitting of the optimized collective variable. The gray landscape represents the density of all simulated paths, with the exception of replicas 4 and 11. All snapshots for which committors were computed are represented by dots colored according to the corresponding committor value. The inset provides zooms in on the approximate reactive path region. The pink dashed line indicates the optimal position of the transition state, and the pink stripe, ranging from -0.25 to 0.25, has been introduced as an extension of the transition state to enable the calculation of the distribution of committor values in the transition state.

To gain a better understanding of the physical processes that underlie the transition of the MscL from the closed state to the S1 state, we divided the optimized reaction coordinate into two uncorrelated parts as detailed in Section 2.4. A 2D space was constructed upon which a density map was plotted (see Figure 3 c). It is noteworthy that all replicas exhibit a comparable behavior, initially moving along the Y-axis and subsequently presumably along the X-axis (the individual trajectories plotted on the same landscape are illustrated in Figures S5 a-n). The physical meaning of these two axes will be elucidated in the subsequent section.

The efficacy of the optimized reaction coordinate in localizing the transition state enabled the calculation of the free energy profile along this reaction coordinate. For this purpose, we employed the umbrella sampling approach (Figure 4 a). The estimated height of the barrier is 10 kJ/mol. The peak of the barrier, which is by definition a transition state, is approximately projected onto the zero value of the reaction coordinate. The closed state basin under the applied tension lies within an interval of -17 to -5.4 (Figure 4 b) and is defined as a region containing all points to the left of the barrier whose energy does not exceed the minimum energy by more than 1 RT. The S1 state is similarly defined as a basin to the right of the barrier with a reaction coordinate falling between 6.7 and 9.8. The free energy difference between the closed and S1 states is proportional to the logarithm of the ratio of populations of these states. Umbrella sampling provides a histogram of the population along the collective variable. To obtain the population of a state, we integrated the histogram with respect to the collective variable over that state. The resulting free energy difference between the closed and S1 states is estimated to be 5 kJ/mol. As anticipated, the S1 state is more favorable than the closed state under applied tension; otherwise, channel opening would be unlikely. To demonstrate the convergence of the free energy profile, it was calculated on multiple occasions throughout the equilibration process. It is notable that the transition state exhibited only a slight decrease in free energy during the equilibration process, indicating that the optimized collective variable is nearly optimal. The final step in validating the quality of the committor-based reaction coordinate is to ascertain whether the free energy profile is capable of reproducing the average transition time between the closed state under tension and the S1 state. The "true" average transition time has been determined to be 133 ± 13 ns (Figure 4 c). By employing the free energy barrier height, we were able to ascertain that k_TST_ is 0.02 s⁻¹ and the corresponding average transition time is 50 ns. Despite the fact that the estimated average transition time is lower than the corresponding "true" value, the agreement is nevertheless satisfactory. This finding indicates that the computed free energy profile is, for the most part, accurate and the committor-based reaction coordinate can be regarded as optimal.

**Figure 4.**
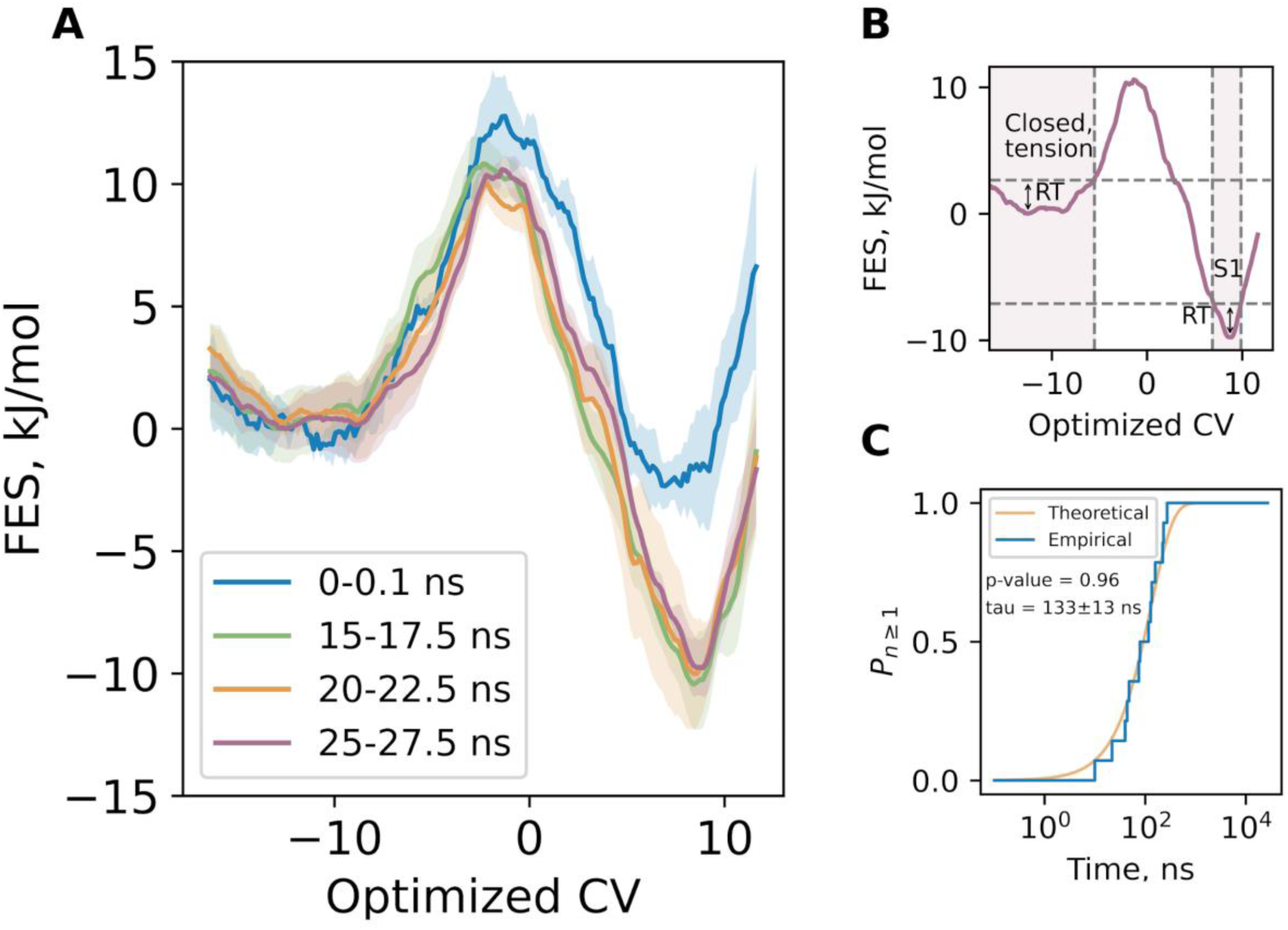
Thermodynamics and kinetics characteristics of the transition from the closed to S1 state. **A.** The free energy profiles along the optimized collective variable at various stages of the equilibration process, accompanied by a standard error of the mean. **B.** The definition of the closed state under the applied tension and the S1 state, based on the RT level. **C.** The empirical (blue) and theoretical (yellow) cumulative distribution function of the transition times from the closed state under tension to the S1 state. The two-sample Kolmogorov–Smirnov test confirms with p-value of 0.96 that the empirical times belong to the exponential distribution with an average time of 133 ± 13 ns.

### The main factors contributing to the reaction coordinate

An analysis of the closed state of MscL reveals that it is stabilized by multiple contact sites (refer to Section S1). Among these sites, the "periplasmic" and "cytoplasmic" sites are disrupted by tension; however, only the disruption of two "cytoplasmic" sites, formed with the participation of the B-chain, facilitates the transition between the closed and S1 states.

During the transition, tensile forces induce substantial modifications to the closed state structure (Figures 5a-b, S7b), resulting in the near-disruption of the ’periplasmic’ site (e.g., contacts T40b:S74b, T35b:N78b) and substantial weakening of the ’cytoplasmic’ site between chains A and B (e.g., contacts A18a-I23b, T25a-T28b).The cytoplasmic site between chains B and C remains intact. The metastable state (CV=-10) resulting from these forces is referred to as the closed state under tension (Figure 4b, left basin). This metastable state is also referred to in literature as the funnel-shaped structure, as it appears much more open from the periplasmic side of the membrane. A two-dimensional representation of the optimized reaction coordinate (Figure 3c) reveals that the path from the closed state at zero tension towards the funnel-shaped structure predominantly lies along the Y axis. As the break of the "periplasmic" site does not contribute to the reaction coordinate at all, the movement along the Y axis is therefore connected with the weakening of the "cytoplasmic" site between chains A and B. The present study indicates that this event marks the initial pivotal moment in the transition of MscL from the closed to the S1 state.

**Figure 5.**
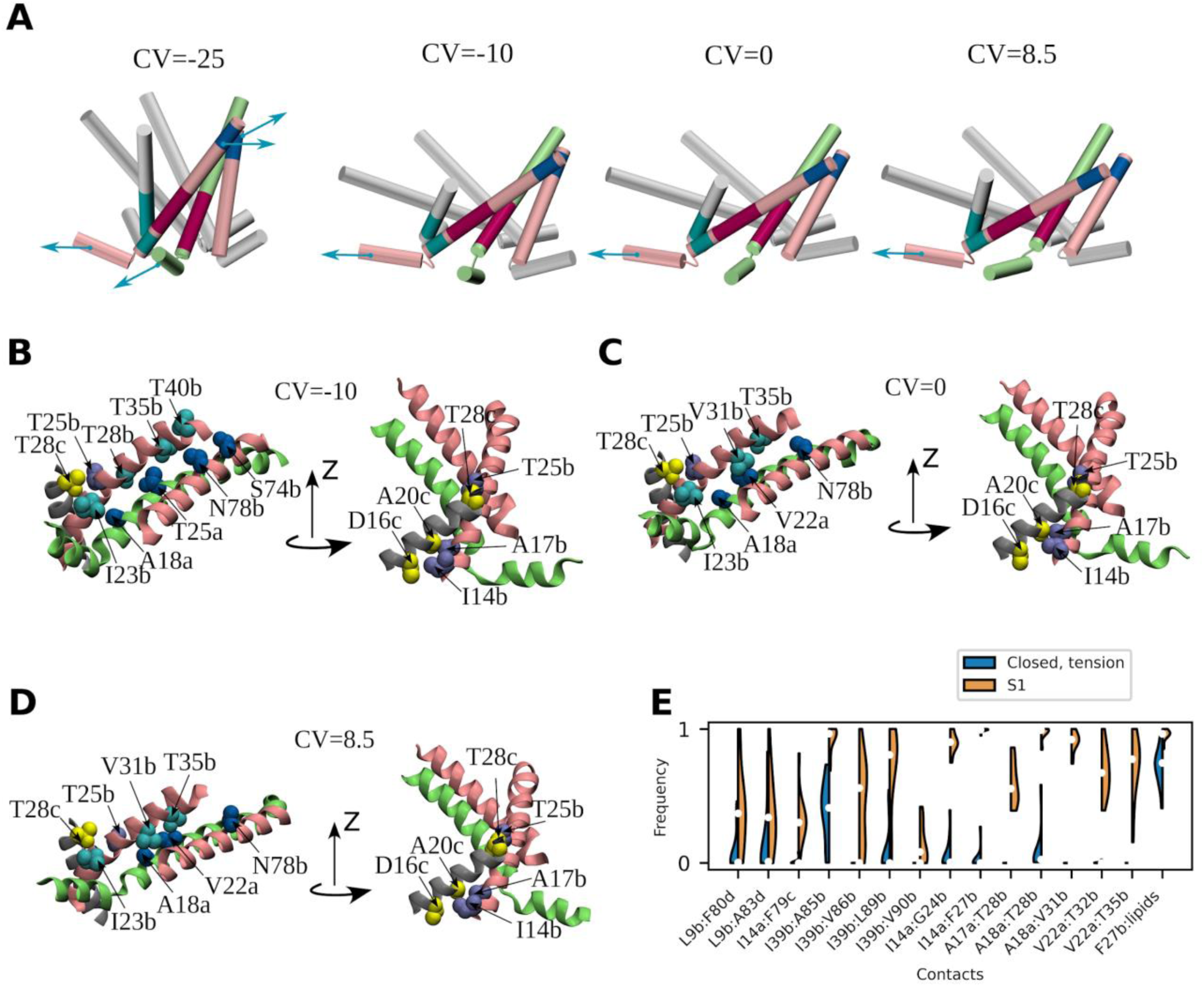
MscL transition from the closed state to the S1 state performed by chain B. **A.** Individual snapshots capture the following states: closed state at zero tension, closed state under tension (“funnel-shaped structure”), transition state, and S1 state. Here and in panels b-d, chains B and A are shown in pink and lime, respectively. For chains A, C, D, and E, only the TM1 and N-terminal helices are shown. The approximate direction of the tensile forces, acting on the ribs involved in the conformational transition is indicated by blue arrows. Colored sections represent parts of the three contact sites most affected during the transition: blue - first contact site between TM1 and TM2 helices of chain B, purple - second site between TM1 helices of chains A and B, cyan - second site between TM1 helices of chains B and C. **B.** View of selected contacts from the above locations in the closed state under tension, **C.** transition state, and **D.** S1 state. **E.** Frequency of occurrence of the contacts typical of the S1 state in the closed state under tension and in the S1 state. To plot the frequencies, two residues were considered to be in contact if the minimum distance between any two heavy atoms of the corresponding residues was less than 4.5 Å. White circles represent the medians, black thick lines run from the first to the third quartile of the distributions.

As illustrated in Figure 3c, upon crossing the free energy barrier and entering the S1 state, both parts of the reaction coordinate undergo a change, with the contribution along the X axis becoming more pronounced. A comparison of the closed state under tension, the transition state, and the S1 states provides insight into the physical process underlying this part of the reaction coordinate.

In the transition state, some contacts belonging to the ‘cytoplasmic’ site between chains B and C (e.g. I14b:D16c, T25b:T28c, A17b:A20c) are weakened, whereas the analogous site of contacts between chains A and B is even more disrupted (Figures 5c, S7c). To illustrate, the distance between residues A18a and I23b increases from 4.4±0.1Å in the closed state at tension to 6.2±0.1Å in the transition state, thereby preventing the two residues from forming a contact. Conversely, A18a and V31b, as well as V22a and T35b, move closer to each other and begin to form contacts. These coupled changes suggest that the TM1 helix of chain B moves with respect to the rib, formed by the TM1 helix of chain A and TM2 helix of chain B. The sole factor that can account for the observed movement of the TM1 helix of chain B is pulling of this chain by means of the N-terminal tension sensor (as shown in Figure 5 and in the movie S9). It is important to note that the TM1 helix of chain B forms a rib with the TM2 helix of chain C. Consequently, all forces applied to the N-terminal helix of chain B are transmitted not only to the TM1 helix of the same chain but also to the entire rib, and therefore the entire rib is subjected to a pulling force.

As illustrated in Figure S7c, the transition state is also characterized by a reduction in the number of contacts between the acyl chains and the lipid-binding pocket. This observation is of particular importance because, in our simulations, MscL was unable to cross the barrier and enter the S1 state without the removal of lipids from this pocket. The loss of contacts between the lipids and the residues forming the binding pocket correlates with committor values and contributes together with the B-chain pulling to the part of the reaction coordinate, located along the X-axis in Figure 3c. The function of lipids in MscL gating will be addressed in greater detail in Section S2.

In accordance with the B chain displacement, the S1 state is characterized by the complete loss of the majority of stable contacts that were present in the closed state under zero tension (Figures 5d, S7d). Conversely, a set of novel contacts, which serve to fix the new position of the B chain, emerge, including A18a:V31b and V22a:T35b (Figures 5d-e). It is noteworthy that the interactions between residues V22a and T35b have been previously studied. Through disulfide trapping experiments^10^, it was determined that these residues are in proximity in the open state of MscL, a result that is corroborated by the present data.

The S1 state is distinguished by its conspicuous pore size, which renders it conductive (see section S3). Another anticipated attribute of the S1 state is the substantial weakening and even loss of some contacts between the TM1 helices of chains B and C (Figures 5d, S7d). This observation suggests that chain C may play a pivotal role in the transition to the S2 state.

### Cooperativity between transitions of different chains

Upon examination of the "funnel-shaped structure" (Figure 5a), the movements of all ribs appear comparable, suggesting a degree of cooperativity between them. However, given the observation that all ribs possess the same tension sensors and are subjected to analogous tension forces, the presence of cooperativity becomes less evident. Indeed, the observed movements of the ribs do not correlate with the computed committor values for the transition of chain B. Furthermore, the positions of the ribs differ significantly between replicas in the S1 state, indicating that the transitions of the five MscL chains can be considered as occurring independently in the initial approximation. However, the observation that the movement of chain B affects the ’cytoplasmic’ site of contacts between chains B and C indicates the potential for limited cooperativity between the conformational transitions of different chains.

As previously discussed in Section 3.2, in order to pull chain B, it is necessary to break the contacts between chains B and C. However, it should be noted that breaking some contacts between other chains also correlates with committor values. The locations of the broken contacts are illustrated in Figures 6a and 6b. If the contacts between chains A and B (illustrated in red) are not considered, the greatest loss of contact occurs between the TM1 helices of chains B and C (depicted in blue). The next largest set of contacts that are weakened by the transition of chain B is highlighted in light blue and located between the TM1 helices of chains D and E. This site is particularly vulnerable due to the positioning of chain D, which is situated opposite chain B within the pentagon. As one side of the pentagon undergoes an increase, the opposite side attempts to react in order to maintain the overall symmetry of the structure. The remaining pairs of chains, specifically C and D and E and A, are only minimally affected by the transition of the B chain. It can thus be proposed that the TM1 helix of C chain and, to a lesser extent, the TM1 helix of E chain have a higher probability of being pulled following the B chain than TM1 helices of chains A and D. This hypothesis is consistent with the results of our simulations, which quantify the frequency of each chain undergoing a transition towards the S2 state. (Figure 6c). In over half of the cases, the subsequent chain to respond to applied tension and perform the transition towards the open state was chain C. However, in slightly less than half of the cases, other chains were pulled first, indicating that stochastic processes exert a considerable influence on the sequence of chain openings.

**Figure 6.**
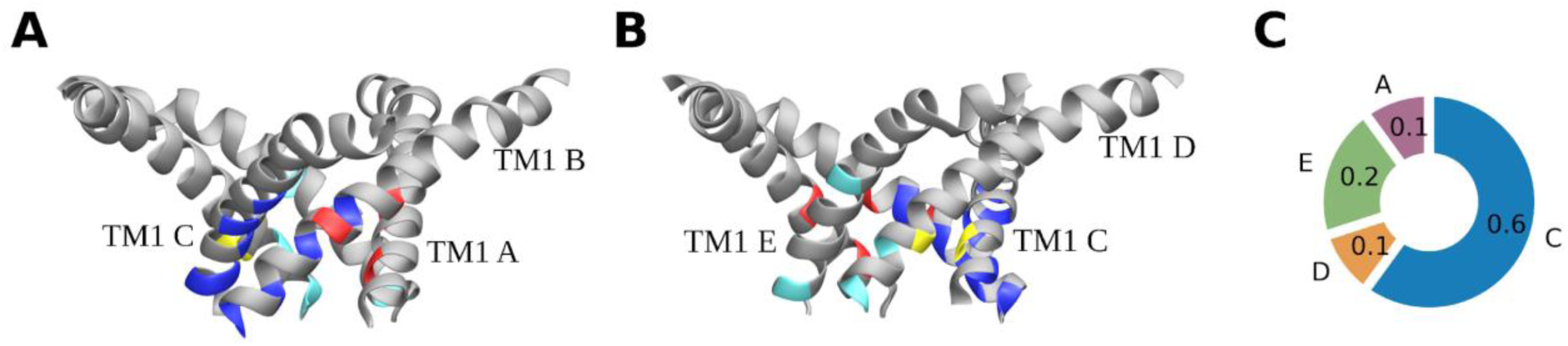
Cooperativity effect between conformational transitions of different chains of MscL. **A.** The representation portrays the locations of contacts between pairs of adjacent chains. The weakening of these contacts correlates, to a certain extent, with the committor values that have been calculated for the transition of chain B. For the sake of simplicity, the illustration depicts only the TM1 helices of all chains. The contacts between chains A and B are highlighted in red, between chains B and C in dark blue, between chains C and D in yellow, and between chains D and E in light blue. **B.** The figure shows the same structure as in figure **A.**, but from a different angle. **C.** The frequency with which each chain undergoes a conformational transition following the conformational transition of chain B.

## DISCUSSION

In the present study, we employed committor values and a nonlinear regression model to construct a reaction coordinate as an optimal linear combination of features selected from a flexible set. A committor function is, by its very nature, an ideal reaction coordinate. However, it is not a suitable standalone tool for gaining mechanistic insight into a given process, as it does not depend in a transparent way on the physical parameters of the system. It is therefore intuitive to employ methodologies that establish a relationship between the committor function and the characteristics of the system. Several methods for addressing this issue have been proposed. Ma and Dinner integrated a neural network and a genetic algorithm to evaluate a comprehensive set of features, identify an optimal subset, and derive an analytical expression for the reaction coordinate^46^. Hummer and Best did not calculate committor values directly; rather, they optimized the p(TP|r) function, which represents the probability of being on a transition path (TP) given a specific value of the reaction coordinate r^47^. As demonstrated, this function is analytically related to the committor for the limit of diffusive dynamics. Peters and Trout implemented an aimless shooting algorithm, which is a variation of the transition path sampling method^48,49^. This results in a binary committor outcome (0 or 1) for each shooting point. By employing a sigmoid function to correlate committor values with a reaction coordinate and characterizing a trial reaction coordinate as a linear combination of a flexible set of features, they maximized the likelihood of obtaining the observed committor outcome. Recently, this approach was modified to account for the continuous nature of the committor function^50^. The computation of committor values and employment of a deep learning algorithm were successfully implemented to iteratively improve the sampling of the transition state^51^. The corresponding nonlinear estimate of the optimal reaction coordinate was then found using symbolic regression. In a most recent paper, the path collective variable was optimized by performing a kernel ridge regression of the committor values using a flexible basic set of features^52^.

The present methodology is relatively uncomplicated and is in accordance with the ideas put forth in these studies. A sigmoid function was employed to approximate a committor function, and a trial reaction coordinate was introduced as a linear combination of a flexible set of features. The committor function was treated as a continuous variable. In lieu of deep learning algorithms, we implemented a straightforward nonlinear regression model. To circumvent overfitting, the model was augmented with optimization of the adjusted R² coefficient via a Monte Carlo procedure. Our findings indicate that even this relatively simple procedure, which does not necessitate sophisticated expertise or substantial computational resources, can yield meaningful outcomes in intricate real-world systems.

The optimized collective variable constructed in this study provides an adequate fit of the committor values with a sigmoid curve, effectively discriminating between basins and the transition state. This is corroborated by the free energy profile, which exhibits a clear peak at nearly zero (the value of the collective variable, by construction, representing a transition state) and shows a relatively rapid convergence to a finite free energy profile with a minimal decrease in barrier height at equilibrium. The height of the barrier is consistent with the average transition time computed from the trajectories. Collectively, these observations indicate that the optimized collective variable is an adequate approximation of the ideal reaction coordinate.

In the present study, we employed all-atom unbiased molecular dynamics and the GOF L17A, V21A mutant to simulate the MscL under the tension of 30 mN/m. This combination of conditions allowed for the observation of multiple native pathways traversing the barrier from the closed toward the open state within a reasonable time frame. The fact that the pathways are absolutely unbiased facilitates the understanding of the channel opening mechanism.

One of the most significant findings of this study is that the conformational transitions of distinct MscL chains are highly improbable to occur concurrently. This phenomenon can be explained from the standpoint of kinetics. It is more probable that a system will overcome five low free energy barriers than one barrier that is five times higher. Furthermore, the sequential transitions may be facilitated by the cooperative effect, such that each subsequent barrier may, in fact, be lower than the previous one. The stochastic nature of the tensile force and the structural independence of the ten tension sensors also favor the hypothesis of sequential openings of the distinct chains. The resulting states with one and two open chains are designated as S1 and S2, respectively. It is noteworthy that the S1 state of another MscL GOF mutant, V21D, has been previously reported in the literature (Figure 4f in reference 11), although it was not discussed. This state was reached under a tension of 50 mN/m by 350 ns of simulations, which is in good agreement with our study. Asymmetric intermediates in the MscL opening pathway were also observed in experiments^31,32^, and it was noted that they need to be accounted for in models of MscL gating. However, the lack of sufficient experimental data precludes a comparison of these asymmetric intermediates with the S1, S2, and other asymmetric states introduced in this study.

In this study, we also investigated whether there is a cooperative effect between the opening of the distinct chains. No strict order was observed, which is indicative of the limited stochasicity of the choice of chains. However, if the chain "i" is already opened, the "i+1" chain, situated clockwise when viewed from the periplasmic side of the channel, has an elevated probability of being opened subsequently. To ascertain whether the chain "i" also exhibits a positive cooperative effect on chain "i+3," larger statistical samples are required.

It is noteworthy that our simulations revealed a multitude of transitions from the closed to the S1 state. However, we did not observe any backward transitions, with the exception of brief excursions between the basins when the system was situated in the vicinity of the transition state region. The lack of transitions from the S1 to the closed state can be attributed to the observation that the transition “S1→S2” appears to have a lower barrier than the transition “S1→Closed”. Although the precise height of the barrier for the “S1 →S2” transition remains unknown, it is reasonable to assume that it cannot exceed the barrier for the “Closed→ S1” transition due to the cooperative effect. This assumption is corroborated by our observation that the “S1→S2” transition occurs more rapidly than the “Closed→S1” transition. The kinetically guided transition into the open state through several sequential stages can act as a ratchet, effectively preventing the channel from spontaneously closing when the big pore is critical for cell survival. This mechanism provides an additional biological explanation for why sequential opening of the different chains can confer evolutionary advantages.

The results of our simulations suggest that two principal contact sites are responsible for maintaining the closed state of the MscL. The “periplasmic” site is situated between the TM1 and TM2 helices of the same chain, in close proximity to the periplasmic side of the membrane. It is structurally linked to the periplasmic tension sensor (L72). This site is almost entirely disassembled when the system reaches the basin of the closed state under tension, and its disassembly renders the TM1 helix of the same chain susceptible to further tensile forces applied to the N-terminal tension sensor. The process of destroying this site of contacts and reaching the basin of the closed state under tension has been previously reported in the literature as an "asymmetric expansion into a funnel-like structure"^11^. In this context, the term "asymmetric" is used to describe a phenomenon in which the part of the channel located closer to the periplasmic side of the membrane undergoes expansion, whereas the part of the protein closer to the intracellular side of the membrane remains tightly packed. In the closed-state basin under tension, we observed the presence of funnel-shaped structures that were almost perfectly shaped. However, in some replicas, we observed structures with varying degrees of opening of individual chains. We believe that this observation should not be overlooked, as the non-simultaneous opening of individual chains is an intrinsic feature of MscL. The “cytoplasmic” site is situated between the TM1 helices of the adjacent chains in close proximity to the cytoplasmic side of the membrane and in association with the intracellular N-terminal tension sensor. In the wild-type protein, this site comprises L17 and V21 residues, which form a well-described “vapor-lock”^16^. Both residues are mutated in the current study; nevertheless, the remainder of this site still preserves and contributes significantly to the stability of the closed state under tension. This is evidenced by the fact that the breaking of this site correlates with the computed committor values.

The pivotal function of the L17 and V21 residues in maintaining the channel in a closed state can be attributed to two factors, both of which contribute to an increase in the height of the free energy barrier for MscL gating. Firstly, the L17 and L21 residues are responsible for maintaining the stability of the contact site between the TM1 helices. Secondly, these residues constitute the hydrophobic gate of the channel. As previously proposed in reference 11, due to their hydrophobic nature, they resist the wetting of the pore. The same paper reports that in 20 μs simulations under 50 mN/m tension, the wild-type MscL expanded into a funnel-like structure with trans-membrane helices bent by nearly 70°. However, it exhibited only intermittent, limited wetting of the hydrophobic gate. Conversely, in the most severe GOF mutants, such as V21D, the gate was wetted even in the absence of applied tension. In our simulations of the L17A, V21A GOF mutant, the gate became wetted in the closed state only when tension was applied, resulting in a limited leaking conductance of 0.3 ± 0.1 nS. The subunit-averaged bend angle of the TM1 helix, as defined in the referenced paper, did not exceed 60° in this state. This confirms that the N-terminal parts of the TM1 helices are slightly displaced from each other and that the gate is slightly open.

While the L17 and V21 residues are of particular importance for the prevention of spontaneous MscL gating, it seems reasonable to assume that their influence does not extend beyond the hydrophobic gate and the "cytoplasmic" contact site. Therefore, the opening process of the wild-type MscL is likely to follow a comparable mechanism to that of the mutant protein, and our findings can be employed to elucidate the wild-type protein gating. This assumption is corroborated by preceding coarse-grained and all-atom simulations of both the wild-type and GOF mutants of MscL, which did not reveal any substantial discrepancies in the gating process^11, 12^.

A further significant outcome of our investigation is the discovery that the acyl chains of lipids occupy the binding pocket formed by I82, V86, and V22 residues in the closed state, thereby impeding MscL gating. The process of delipidation of this pocket is found to correlate well with the committor values computed for the MscL transition between the closed and the S1 states. It seems reasonable to posit that the delipidation of the other pockets will contribute to the transitions of the other chains, leading to the states S2, S3, and so on. It is noteworthy that the significance of delipidation of this pocket was recently reported for the first time^7,19,53^. A series of studies have demonstrated that introducing bulky modifications of the L89 residue restricts lipid-chain access to the binding pocket, resulting in a notable overall reduction in MscL’s pressure activation threshold and stabilization of the conductive state.

The validation of the S1 state structure is complicated by the fact that it is a very short-lived stage and therefore unlikely to be observed in experiments. However, we may conceptualize S1 as a partially open state, in which only one chain undergoes conformational transitions, and compare the interactions observed in our simulations for this single chain with those derived from experiments investigating fully open and expanded conformations.

The sole experimentally resolved expanded structure of MscL is derived from M. acetivorans^14^. This expanded state is not entirely open; rather, it is trapped in an intermediate state, as can be concluded based on the in-plane protein area^14,54^. We performed a structural alignment between the "expanded" fragment of the S1 state of MtMscL and the expanded state of MaMscL, utilising a sequence alignment between these proteins (Figure 1c). A structural alignment was conducted using the protein sequences that constitute two adjacent ribs of MtMscL, which are separated in the S1 state: I14a-I38a, I14b-I38b, L69b-L89b, and L69c-L89c. The aligned structures are shown in Figure 1d. Firstly, we focused on the contact between residues V22a and T35b of MtMscL, as this was observed not only in the S1 state but also documented by disulfide trapping experiments. The homologous contact in the expanded state of MaMscL was subsequently identified as occurring between residues I24 and V37′ (here, the apostrophe mark designates another chain).

Two further pairs of contacts in MaMscL, for which we endeavoured to identify homologues within the MtMscL structures, are A20:A22′, V37′:N77′ in the closed state and A20:S29′, V37′:I80′ in the expanded state^14^. The homologous contacts in the closed state of MtMscL are A18a:A20b and T35b:N78b, which were identified as the stable contacts at zero tension (Figure S7a). In the S1 state the homologous contacts should be A18a:F27b and T35b:L81b. Indeed, for A18a, we observed a highly stable contact with T28b (Figure 5e) and a less significant contact with F27b, which did not contribute to the optimized collective variable. It is regrettable that the V37′:I80′ contact cannot be directly matched to the MtMscL contacts, as it does not truly exist in the expanded state of MaMscL, as evidenced by the 4Y7J entry in the PDB. It is important to note, however, that in the S1 state described in the current study, a weak contact was observed between the T35b and I82b residues. As indicated by the provided alignment, this corresponds to the V37′:I81′ contact in MaMscL, which is indeed present in the expanded state. Therefore, the aligned structures exhibit an appropriate fit, suggesting that the S1 state of MscL identified in the current study represents an intermediate form that ultimately leads to the expanded state.

The open state of MscL can be partially characterized by disulfide trapping experiments conducted on the E. coli protein. The experimental data indicates the presence of five contacts: V23:I96, C26:I92, I32:N81, I24:V37, and A20:L36^10,30^. As evidenced by the alignment of MscL sequences^19^, these contacts are homologous to V21:V91, G24:V86, L30:A75, V22:T35, and A18:F34 in the MtMscL structure. L30:A75 is one of the contacts that maintains the bonding between the TM1 and TM2 helices, which belong to different chains but are in the same rib. This is why it is present in both the closed state, as reported by the crystal structure (PDB ID 2OAR), and the S1 state, as can be observed in our simulations. The remaining pairs of residues are unable to form the specified contacts in the closed state due to their considerable distance from one another. However, the S1 state provides a greater potential for these interactions to occur. The V22a:T35b interaction is not only highly stable in the S1 state, but its formation also contributes to the optimized collective variable. The F34b residue is approaching the A18a residue as a result of the conformational transition from the closed to the S1 state. While the contact has not yet been formed, it appears that pulling the TM1 helix of chain B one additional turn may facilitate its establishment. In our preliminary unbiased simulations under the applied tension of 30 mN/m, we observed that the chain B was pulled further and that the A18a:F34b contact was formed. Notably, this process did not affect the L30:A75 and V22a:T35b contacts.

It is plausible that contacts V21b:V91c and G24b:V86c could have been formed in the S1 state if the TM1 helix of chain B and the TM2 helix of chain C had undergone a rotation around their axes. Noteworthy, a clockwise rotation of the TM1 helix and a counterclockwise rotation of the TM2 helix have been previously reported^7,55^. In these studies, the direction of rotation was defined when the MscL channel was viewed from the periplasmic side. In accordance with this consensus, we observed a clockwise rotation of the TM1 helix of chain B in the S1 state. We hypothesized that this process is at least partially guided by an increase in interaction between the hydrophobic side chain of F27b and surrounding lipid tails. Although the direction of the observed rotation is consistent with the previous findings, the magnitude of this motion in our simulations was insufficient to form the aforementioned contacts. Additionally, our simulations indicate a slight counterclockwise rotation of the TM2 helix of chain C in the S1 state. However, the reliability of this observation is questionable, and further investigation is necessary to confirm its validity.

A comparison between the S1 state and both the open and intermediate states provides evidence that the S1 state is not fully open, even with regard to chain B. In order to achieve the final position of this chain, as in the true open state, the rib formed by the TM1 helix of chain B and the TM2 helix of chain C should be pulled one additional helical turn to facilitate the formation of the A18a:F34b contact. Furthermore, the helices should undergo a subsequent rotation (TM1 helix clockwise and TM2 helix counterclockwise), which appears to be a two-gear mechanism. Nevertheless, it is assumed that the S1 state, as well as the closed state under tension, represent the initial genuine metastable states along the pathway of MscL opening. A comprehensive description of these states would facilitate the determination of the structure of the open state and a more profound comprehension of the MscL gating mechanism.

In conclusion, we put forth a model that encapsulates the primary stages of MscL opening, as evidenced by the results of our simulations (Figure 7). For the sake of simplicity, we have depicted the pseudo-channel, comprising only three ribs. The figure illustrates the projection of the ribs onto the membrane plane. In the absence of tension, the adjacent ribs are held together by the “periplasmic” and “cytoplasmic” sites of contact, resulting in an orientation of the ribs that is almost orthogonal to the membrane plane. A junction between two adjacent ribs was defined as a set of two Cα atoms (one from the TM1 helix of the first rib and the other from the TM1 helix of the second rib) such that the distance between these atoms is the minimum among the distances for all possible such sets. In the closed state at zero tension, the junction is formed by Cα atoms of A18 and A20. The Cα atoms of the A20 residues are indicated by blue circles. When tension is applied, tensile forces act on the cytoplasmic and periplasmic sensors, which could cause torques. However, the intracellular part of the channel is rather resistant to stretching due to the very stable intracellular site of contacts. In contrast, the periplasmic part responds readily to tensile forces by stretching. Consequently, MscL attains the "funnel-like" configuration, which represents the closed state under tension. The perfect "funnel-like" structure, exhibiting a uniform degree of opening for all ribs, is illustrated in the figure. In a GOF mutant with an impaired intracellular contact site, both periplasmic and intracellular tension sensors respond to the applied tension at this step. This is evidenced by the substitution of residue A20 for G24 in a junction with A18, which is a consequence of the ribs being pulled one helical turn via the intracellular sensor. The Cα atoms of the G24 residues are indicated by magenta circles. It seems implausible that the wild- type protein would lose the junction between the A18 and A20 residues at this stage. The final step illustrated in the figure represents the transition of MscL into the S1 state. Tensile forces are applied to all tension sensors, but only one responds by pulling the rib one additional helical turn, as this pulling is accompanied by overcoming a free energy barrier. The new junction position is between the Cα atoms of A18 and T28. The Cα atoms of the T28 residues are indicated by yellow circles. As previously discussed, the further pulling of all ribs and the gear-like rotation of their constituent helices should result in the formation of the completely open state.

**Figure 7.**
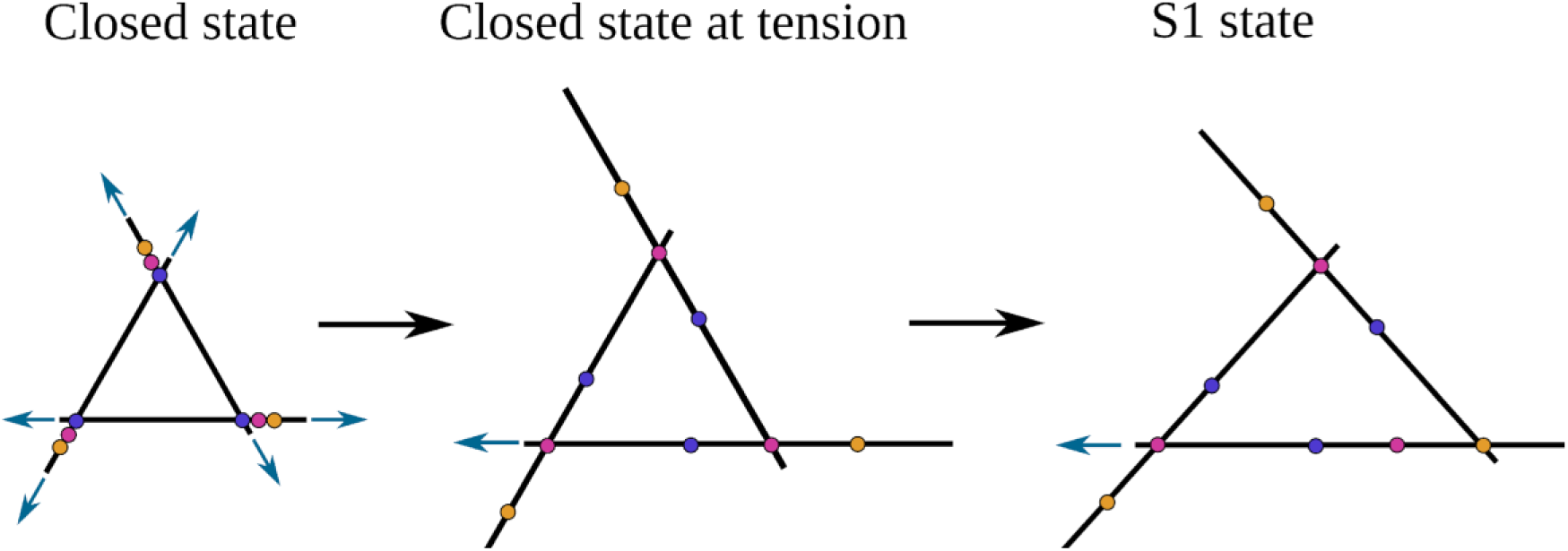
A model of the MscL transition from the closed to the S1 state. In this figure, the pseudo-channel with three chains is considered. The black lines represent the projections of the three ribs onto the membrane plane. Given that the length of the ribs remains constant, the longer projection indicates a reduction in the angle between the rib and the membrane plane. The blue arrows represent the tensile forces acting on the periplasmic and cytoplasmic tension sensors of the channel. Although the tensile forces act on all sensors, in the second and third representations, we have demonstrated only the force that acts on the intracellular sensor of the chain that is pulled. The Cα atoms of the residues A20, G24, and T28 are illustrated with blue, magenta, and yellow circles, respectively. The junction between two adjacent ribs is invariably formed by the A18 residue from one rib (not shown) and one of the residues A20, G24, or T28 from the other rib, contingent on the extent of this latter rib’s displacement.

## Supporting information

Supplementary Material pdb file

Supplementary Material input file

## ASSOCIATED CONTENT

### Data and Software availability statement

The closed state of MscL at zero tension was taken from the PDB with ID 2OAR. (https://www.rcsb.org/structure/2OAR). GROMACS 2020.7 and GROMACS 2022.5 patched with PLUMED 2.8.2 were used to perform all MD simulations (https://www.gromacs.org/ and https://www.plumed.org/). The molecular structures were rendered using the Visual Molecular Dynamics (VMD) program (https://www.ks.uiuc.edu/Research/vmd/). Charts and plots were made using the Python library Matplotlib (https://matplotlib.org/). A "true" average transition time was calculated using the author’s script: https://github.com/valsson-group/time-from-biased-simulations-tools. Scripts for committor analysis and optimzation of the reaction coordinate are available on GitHub: https://github.com/olga-r/MscL.

### Supporting information

Details on the methods, additional description of the MscL conformations, analysis of the role of lipids in the activation of MscL (DOC).

Plumed input file, which is designed to calculate the optimized collective variable (TXT). PDB file containing the structural data for the S1 state (PDB).

Movie illustrating the displacement of chain B during the transition from closed to S1 (AVI).

### Author Contributions

W.K. conceived and supervised the project. O.N.R. performed MD simulations and analyzed the data. O.N.R. wrote the manuscript with comments from W.K.

### Funding Sources

Human Frontier Science Program (HFSP) Early Career Research Grant RGY0085/2021 and Max Planck Society.

## ACKNOWLEDGMENT

Both authors acknowledge the funding from the Human Frontier Science Program (HFSP) Early Career Research Grant RGY0085/2021. We thank Andreas Hartel, Tiago Lopes Marta da Costa and Bert L. de Groot for helpful discussions.

## REFERENCES

1. Kung, C.; Martinac, B.; Sukharev, S. Mechanosensitive channels in microbes. Annu. Rev. Microbiol., 2010, 64, 313–329. DOI: 10.1146/annurev.micro.112408.134106.

2. Moe, P. C.; Levin, G.; Blount, P. Correlating a protein structure with function of a bacterial mechanosensitive channel. J. Biol. Chem., 2000, 275 (40), 31121–31127. DOI: 10.1074/jbc.M002971200.

3. Moe, P.; Blount, P. Assessment of potential stimuli for mechano-dependent gating of MscL: effects of pressure, tension, and lipid headgroups. Biochemistry, 2005, 44, 12239–12244. DOI: 10.1021/bi0509649.

4. Nomura, T.; Cranfield, C. G.; Deplazes, E.; Owen, D. M.; Macmillan, A.; Battle, A. R.; Constantine, M.; Sokabe, M.; Martinac, B. Differential effects of lipids and lyso-lipids on the mechanosensitivity of the mechanosensitive channels MscL and MscS. Proc. Natl. Acad. Sci. U. S. A., 2012, 109 (22), 8770–8775. DOI: 10.1073/pnas.1200051109.

5. Yoo, J.; Cui, Q. Curvature generation and pressure profile modulation in membrane by lysolipids: insights from coarse-grained simulations. Biophys J., 2009, 97 (8), 2267–2276. DOI: 10.1016/j.bpj.2009.07.051

6. Perozo, E.; Kloda, A.; Cortes, D. Physical principles underlying the transduction of bilayer deformation forces during mechanosensitive channel gating. Nat. Struct. Mol. Biol., 2002, 9, 696– 703. DOI: 10.1038/nsb827

7. Kapsalis, C.; Wang, B.; Mkami, H. El.; Pitt, S. J.; Schnell, J. R.; Smith, T. K.; Lippiat, J. D.; Bode, B. E.; Pliotas, C. Allosteric activation of an ion channel triggered by modification of mechanosensitive nano-pockets. Nat. Commun. 2019, 10 (1), 4619. DOI: 10.1038/s41467-019-12591-x.

8. Chang, G.; Spencer, R. H.; Lee, A. T.; Barclay, M. T.; Rees, D. C. Structure of the MscL homolog from Mycobacterium tuberculosis: a gated mechanosensitive ion channel. Science, 1998, 282 (5397), 2220–2226. DOI: 10.1126/science.282.5397.2220.

9. Steinbacher, S.; Bass, R. B.; Strop, P.; Rees, D. C. Structures of the Prokaryotic Mechanosensitive Channels MscL and MscS. Curr. Top. Membr., 2007, 58, 1–24. DOI: 10.1016/S1063-5823(06)58001-9

10. Betanzos, M.; Chiang, C. S.; Guy, H. R.; Sukharev, S. A large iris-like expansion of a mechanosensitive channel protein induced by membrane tension. Nat. Struct. Biol., 2002, 9 (9), 704–710. DOI: 10.1038/nsb828

11. Sharma, A.; Anishkin, A.; Sukharev, S.; Vanegas, J. M. Tight hydrophobic core and flexible helices yield MscL with a high tension gating threshold and a membrane area mechanical strain buffer. *Front. Chem. (Lausanne*, Switz.), 2023, 11, 1159032. DOI: 10.3389/fchem.2023.1159032

12. Yefimov, S.; Van der Giessen, E.; Onck, P. R.; Marrink, S. J. Mechanosensitive membrane channels in action. Biophys J., 2008, 94 (8), 2994–3002. DOI: 10.1529/biophysj.107.119966

13. Rajeshwar, T. R.; Anishkin, A.; Sukharev, S.; Vanegas, J. M. Mechanical Activation of MscL Revealed by a Locally Distributed Tension Molecular Dynamics Approach. Biophys J., 2021, 120 (2), 232–242. DOI: 10.1016/j.bpj.2020.11.2274

14. Li, J.; Guo, J.; Ou, X.; Zhang, M.; Li, Y.; Liu, Z. Mechanical coupling of the multiple structural elements of the large-conductance mechanosensitive channel during expansion. Proc. Natl. Acad. Sci. U. S. A., 2015, 112 (34), 10726–10731. DOI: 10.1073/pnas.1503202112

15. Corry, B.; Hurst, A. C.; Pal, P.; Nomura, T.; Rigby, P.; Martinac, B. An improved open- channel structure of MscL determined from FRET confocal microscopy and simulation. J. Gen. Physiol., 2010, 136 (4), 483–494. DOI: 10.1085/jgp.200910376.

16. Anishkin, A.; Akitake, B.; Kamaraju, K.; Chiang, C. S.; Sukharev, S. Hydration properties of mechanosensitive channel pores define the energetics of gating. J. Phys.:Condens. Matter, 2010, 22 (45), 454120. DOI: 10.1088/0953-8984/22/45/454120

17. Bavi, N.; Cortes, D. M.; Cox, C. D.; Rohde, P. R.; Liu, W.; Deitmer, J. W.; Bavi, O.; Strop, P.; Hill, A. P.; Rees, D.; Corry, B.; Perozo, E.; Martinac, B. The role of MscL amphipathic N terminus indicates a blueprint for bilayer-mediated gating of mechanosensitive channels. Nat. Commun., 2016, 7, 11984. DOI: 10.1038/ncomms11984.

18. Sawada, Y.; Nomura, T.; Martinac, B.; Sokabe, M. A novel force transduction pathway from a tension sensor to the gate in the mechano-gating of MscL channel. *Front. Chem. (Lausanne*, Switz.), 2023, 11, 1175443. DOI: 10.3389/fchem.2023.1175443

19. Kapsalis. C; Ma, Y; Bode, B. E.; Pliotas, C. In-Lipid Structure of Pressure-Sensitive Domains Hints Mechanosensitive Channel Functional Diversity. Biophys J., 2020, 119 (2), 448–459. DOI: 10.1016/j.bpj.2020.06.012

20. Robert, X.; Gouet, P. Deciphering key features in protein structures with the new ENDscript server. Nucleic Acids Res. 2014, 42, W320–W324. DOI: 10.1093/nar/gku316.

21. Maurer, J. A.; Dougherty, D. A. A high-throughput screen for MscL channel activity and mutational phenotyping. Biochim Biophys Acta, 2001, 1514(2), 165–169. DOI: 10.1016/s0005-2736(01)00390-x.

22. Ou, X.; Blount, P.; Hoffman, R. J.; Kung, C. One face of a transmembrane helix is crucial in mechanosensitive channel gating. Proc. Natl. Acad. Sci. U. S. A., 1998, 95 (19), 11471–11475. DOI: 10.1073/pnas.95.19.11471

23. Maurer, J. A.; Elmore, D. E.; Lester, H. A.; Dougherty, D. A. Comparing and contrasting Escherichia coli and Mycobacterium tuberculosis mechanosensitive channels (MscL). New gain of function mutations in the loop region. J. Biol. Chem., 2000, 275 (29), 22238–22244. DOI: 10.1074/jbc.M003056200

24. Louhivuori, M.; Risselada, H. J.; van der Giessen, E.; Marrink, S. J. Release of content through mechano-sensitive gates in pressurized liposomes. Proc. Natl. Acad. Sci. U. S. A., 2010, 107, 19856 – 19860. DOI: 10.1073/pnas.1001316107

25. Melo, M. N.; Arnarez, C.; Sikkema, H.; Kumar, N.; Walko, M.; Berendsen, H. J.; Kocer, A.; Marrink, S. J.; Ingólfsson, H. I. High-Throughput Simulations Reveal Membrane-Mediated Effects of Alcohols on MscL Gating. J. Am. Chem. Soc., 2017, 139 (7), 2664–2671. DOI: 10.1021/jacs.6b11091

26. Jeon, J.; Voth, G. A. Gating of the mechanosensitive channel protein MscL: the interplay of membrane and protein. Biophys J., 2008, 94 (9), 3497–3511. DOI: 10.1529/biophysj.107.109850

27. Gullingsrud, J.; Schulten, K. Gating of MscL studied by steered molecular dynamics. Biophys J., 2003, 85 (4), 2087–2099. DOI: 10.1016/S0006-3495(03)74637-2

28. Martinac, A. D.; Bavi, N.; Bavi, O.; Martinac, B. Pulling MscL open via N-terminal and TM1 helices: A computational study towards engineering an MscL nanovalve. PloS One, 2017, 12 (8), e0183822. DOI: 10.1371/journal.pone.0183822

29. Wang, Y.; Liu, Y.; Deberg, H. A.; Nomura, T.; Hoffman, M. T.; Rohde, P. R.; Schulten, K.; Martinac, B.; Selvin, P. R. Single molecule FRET reveals pore size and opening mechanism of a mechano-sensitive ion channel. eLife, 2014, 3:e01834. DOI: 10.7554/eLife.01834.

30. Li, Y.; Wray, R.; Eaton, C.; Blount, P. An open-pore structure of the mechanosensitive channel MscL derived by determining transmembrane domain interactions upon gating. FASEB J., 2009, 23 (7), 2197–2204. DOI: 10.1096/fj.09-129296.

31. Shapovalov, G.; Bass, R.; Rees, D. C.; Lester, H. A. Open-state disulfide crosslinking between Mycobacterium tuberculosis mechanosensitive channel subunits. Biophys J., 2003, 84 (4), 2357–2365. DOI: 10.1016/S0006-3495(03)75041-3.

32. Shapovalov, G.; Lester, H. A. Gating transitions in bacterial ion channels measured at 3 microns resolution. J Gen Physiol., 2004, 124 (2), 151–161. DOI: 10.1085/jgp.200409087.

33. Herrera, N.; Maksaev, G.; Haswell, E. S.; Rees, D. C. Elucidating a role for the cytoplasmic domain in the Mycobacterium tuberculosis mechanosensitive channel of large conductance. Sci. Rep., 2018, 8, 14566. DOI: 10.1038/s41598-018-32536-6

34. Jo, S.; Kim, T.; Im, W. Automated builder and database of protein/membrane complexes for molecular dynamics simulations. PLoS One, 2007, 2 (9), e880. DOI: 10.1371/journal.pone.0000880.

35. Abraham, M. J.; Murtola, T.; Schulz R.; Páll, S.; Smith J. C.; Hess B.; Lindahl E. GROMACS: High performance molecular simulations through multi-level parallelism from laptops to supercomputers, SoftwareX, 2015, 1, 19–25. DOI: 10.1016/j.softx.2015.06.001

36. Páll, S.; Abraham, M. J.; Kutzner, C.; Hess, B.; Lindahl E. Tackling Exascale Software Challenges in Molecular Dynamics Simulations with GROMACS *In S. Markidis & E. Laure (Eds.)*, Solving Software Challenges for Exascale, 2015, 8759, 3–27. DOI: 10.1007/978-3-319-15976-8_1

37. Huang, J.; Rauscher, S.; Nawrocki, G.; Ran, T.; Feig, M.; de Groot, B. L.; Grubmüller, H.; MacKerell, A. D. Jr. CHARMM36m: an improved force field for folded and intrinsically disordered proteins. Nat. Methods, 2017, 4 (1), 71–73. DOI: 10.1038/nmeth.4067.

38. Gromacs Documentation. Release 2020.2, 2020. DOI: 10.5281/zenodo.3773799

39. Humphrey, W.; Dalke, A.; Schulten, K. VMD - Visual Molecular Dynamics, J. Mol. Graphics, 1996, 14, 33–38. DOI: 10.1016/0263-7855(96)00018-5

40. The PLUMED consortium. Promoting transparency and reproducibility in enhanced molecular simulations, Nat. Methods, 2019, 16, 670. DOI: 10.1038/s41592-019-0506-8

41. Tribello, G. A.; Bonomi M.; Branduardi, D.; Camilloni, C.; Bussi, G. PLUMED2: New feathers for an old bird, Comput. Phys. Commun., 2014, 185, 604. DOI: 10.1016/j.cpc.2013.09.018

42. Diez, G.; Nagel, D.; Stock, G. Correlation-Based Feature Selection to Identify Functional Dynamcis in Proteins *J*. Chem. Theory Comput., 2022, 18 (8), 5079–5088. DOI: 10.1021/acs.jctc.2c00337

43. Virtanen, P.; Gommers, R.; Oliphant, T. E.; Haberland, M.; Reddy, T.; Cournapeau, D.; Burovski, E.; Peterson, P.; Weckesser, W.; Bright, J.; van der Walt, S. J.; Brett, M.; Wilson, J.; Millman, K. J.; Mayorov, N.; Nelson, A. R. J.; Jones, E.; Kern, R.; Larson, E.; Carey, C. J.; Polat, İ.; Feng Y.; Moore, E. W.; VanderPlas, J.; Laxalde, D.; Perktold, J.; Cimrman, R.; Henriksen, I.; Quintero E. A. ; Harris, C. R.; Archibald, A. M.; Ribeiro, A. H.; Pedregosa, F.; van Mulbregt, P.; SciPy 1.0 Contributors. SciPy 1.0: Fundamental Algorithms for Scientific Computing in Python. Nat. Methods, 2020, 17 (3), 261-272. DOI: 10.1038/s41592-019-0686-2.

44. Torrie, G. M.; Valleau, J. P. Nonphysical sampling distributions in Monte Carlo free-energy estimation: Umbrella sampling. J. Comput. Phys., 1977, 23 (2), 187–199. DOI: 10.1016/0021-9991(77)90121-8

45. Salvalaglio, M.; Tiwary, P.; Parrinello M. Assessing the Reliability of the Dynamics Reconstructed from Metadynamics. J. Chem. Theory Comput., 2014, 10 (4), 1420–1425. DOI: 10.1021/ct500040r

46. Ma, A.; Dinner, A. R. Automatic method for identifying reaction coordinates in complex systems. J. Phys. Chem. B, 2005, 109, 6769– 6779, DOI: 10.1021/jp045546c

47. Best, R. B.; Hummer, G. Reaction coordinates and rates from transition paths, Proc. Natl. Acad. Sci. U. S. A., 2005, 102, 6732. DOI: 10.1073/pnas.0408098102

48. Peters, B.; Trout, B. L. Obtaining reaction coordinates by likelihood maximization. J. Chem. Phys., 2006, 125, 054108 DOI: 10.1063/1.2234477

49. Bolhuis, P. G.; Chandler, D.; Dellago, C.; Geissler, P. L. Transition path sampling: Throwing ropes over rough mountain passes, in the dark. Annu. Rev. Phys. Chem., 2002, 53, 291– 318, DOI: 10.1146/annurev.physchem.53.082301.113146

50. Mori Y.; Okazaki, K.; Mori, T.; Kim, K.; Matubayasi, N. Learning reaction coordinates via cross-entropy minimization: Application to alanine dipeptide. J. Chem. Phys., 2020, 153 (5), 054115. DOI: 10.1063/5.0009066

51. Jung, H.; Covino, R.; Arjun, A.; Leitold C.; Dellago, C.; Bolhuis, P. G.; Hummer, G. Machine-guided path sampling to discover mechanisms of molecular self-organization. Nat. Comput. Sci., 2023, 3, 334–345. DOI: 10.1038/s43588-023-00428-z

52. France-Lanord A.; Vroylandt, H.; Salanne, M.; Rotenberg, B.; Saitta, A. M., Pietrucci, F. Data-Driven Path Collective Variables. J. Chem. Theory Comput., 2024, 20 (8), 3069–3084. DOI: 10.1021/acs.jctc.4c00123

53. Wang, B.; Lane, B. J.; Kapsalis, C.; Ault, J. R.; Sobott, F.; Mkami, H. El., Calabrese, A. N.; Kalli, A. C.; Pliotas, C. Pocket delipidation induced by membrane tension or modification leads to a structurally analogous mechanosensitive channel state. Structure, 2022, 30 (4), 608–622.e5. DOI: 10.1016/j.str.2021.12.004

54. Anishkin, A.; Chiang, C.-S.; Sukharev, S. Gain-of-function Mutations Reveal Expanded Intermediate States and a Sequential Action of Two Gates in MscL. J. Gen. Physiol, 2005, 125 (2), 155–170. DOI: 10.1085/jgp.200409118

55. Iscla, I.; Levin, G.; Wray, R.; Reynolds, R.; Blount, P. Defining the physical gate of a mechanosensitive channel, MscL, by engineering metal-binding sites. Biophys J., 2004, 87 (5), 3172–3180. DOI: 10.1529/biophysj.104.049833

