## Supplementary Material file for "Asymmetric nature of MscL opening revealed by molecular dynamics simulations"

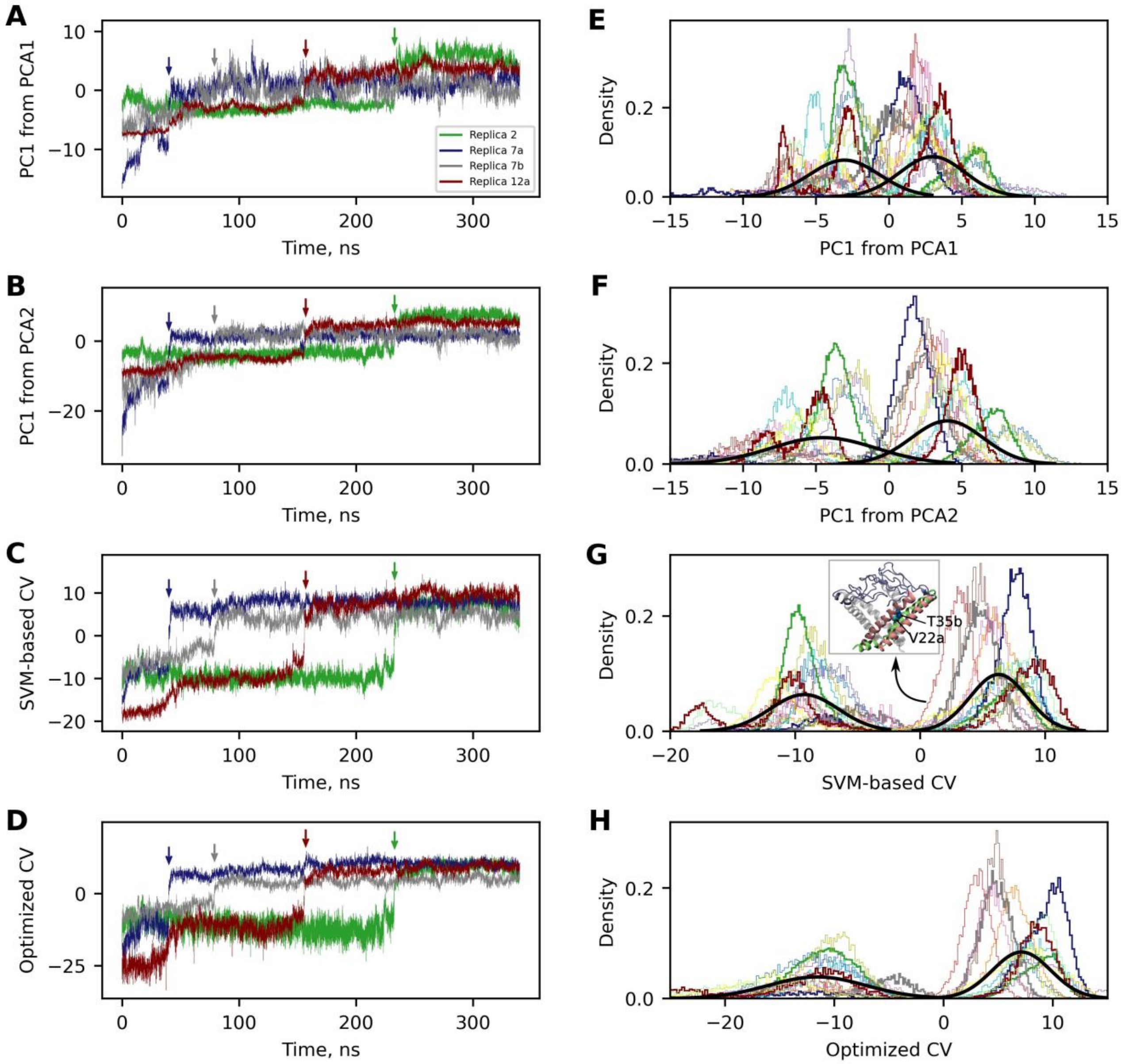


**Figure S1.** Projections of the simulated paths onto the different collective variables: **A** and **E.** PC1 from PCA, applied to stable contacts at zero tension (termed PCA1); **B** and **F.** PC1 from PCA, applied to a large set of contacts that included protein residues and lipids (termed PCA2); **C** and **G.** SVM-based collective variable, derived as a linear combination of the contact and distance SVM-based collective variables; **D** and **H.** Collective variable optimized using computed commitors. **A-D.** Time series of 4 selected paths (the same paths are highlighted with bold in figures **E-H**), arrows point to the events of transition from the closed to S1 state. **E-H.**

Densities of configurations visited by paths, along the collective variables. Black bold Gaussians show the result of applying a Gaussian mixture model to pooled data from all paths. Inset in panel **G** shows the S1 state for Replica 4 (red path). The conformational transition was undertaken by chain B, which is shown in pink. The adjacent TM1 helix of chain A is indicated in lime. Although the protein is in the S1 state, which is confirmed for example by formation of the critical contact V22a-T35b, its conformation appears to be relatively closed, as the flexible loops (ice-blue) are stuck together at the top of the channel.

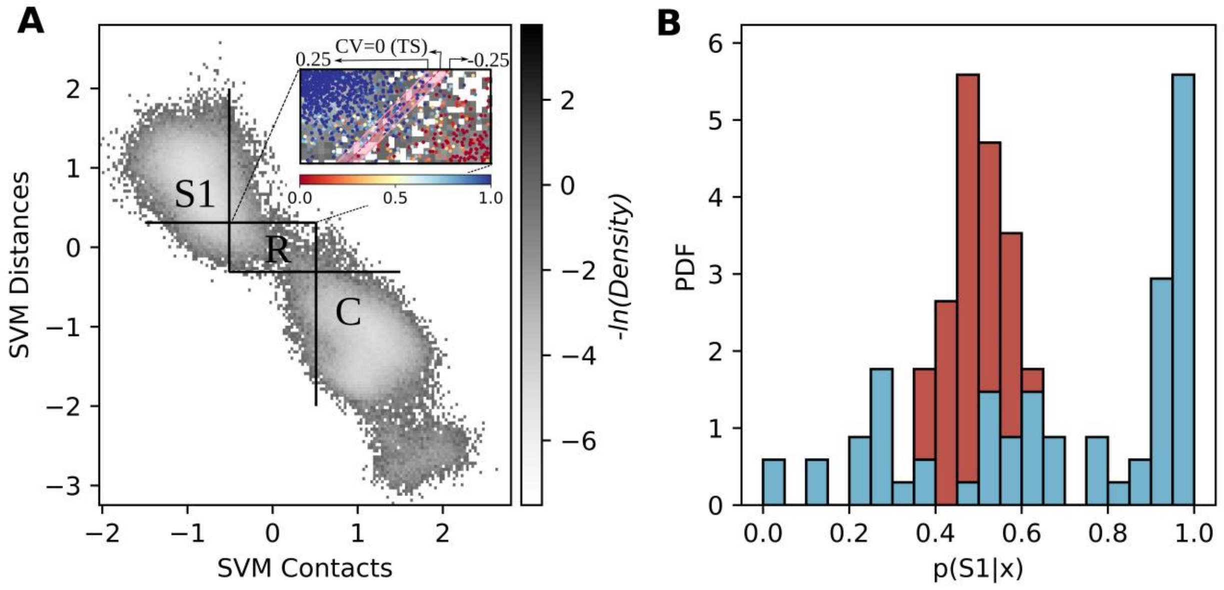


**Figure S2.** Methodology of the committor analysis. **A.** The gray landscape represents a density of all simulated paths, except replica 4 and replica 11, projected onto the space formed by the SVM-based collective variables. The regions approximating the closed (C) and S1 states and the reactive path region (R) are shown. The snapshots for committor computations were selected from the reactive path region. The inset shows all snapshots colored by the corresponding committor value. The pink dashed line indicates the optimal position of the transition state on the SVM-based collective variable. The pink stripe was introduced as an extension of the transition state to allow calculating the distribution of committor values in the transition state. **B.** The distribution of committor values in the transition state, defined in the inset of panel a), is shown in blue. A red histogram is given for comparison to show the approximate appearance of the "ideal" committor distribution. It is obtained by taking a sample of the same size as used for the blue histogram from the Gaussian distribution (mean=0.5, $\sigma$=0.07).

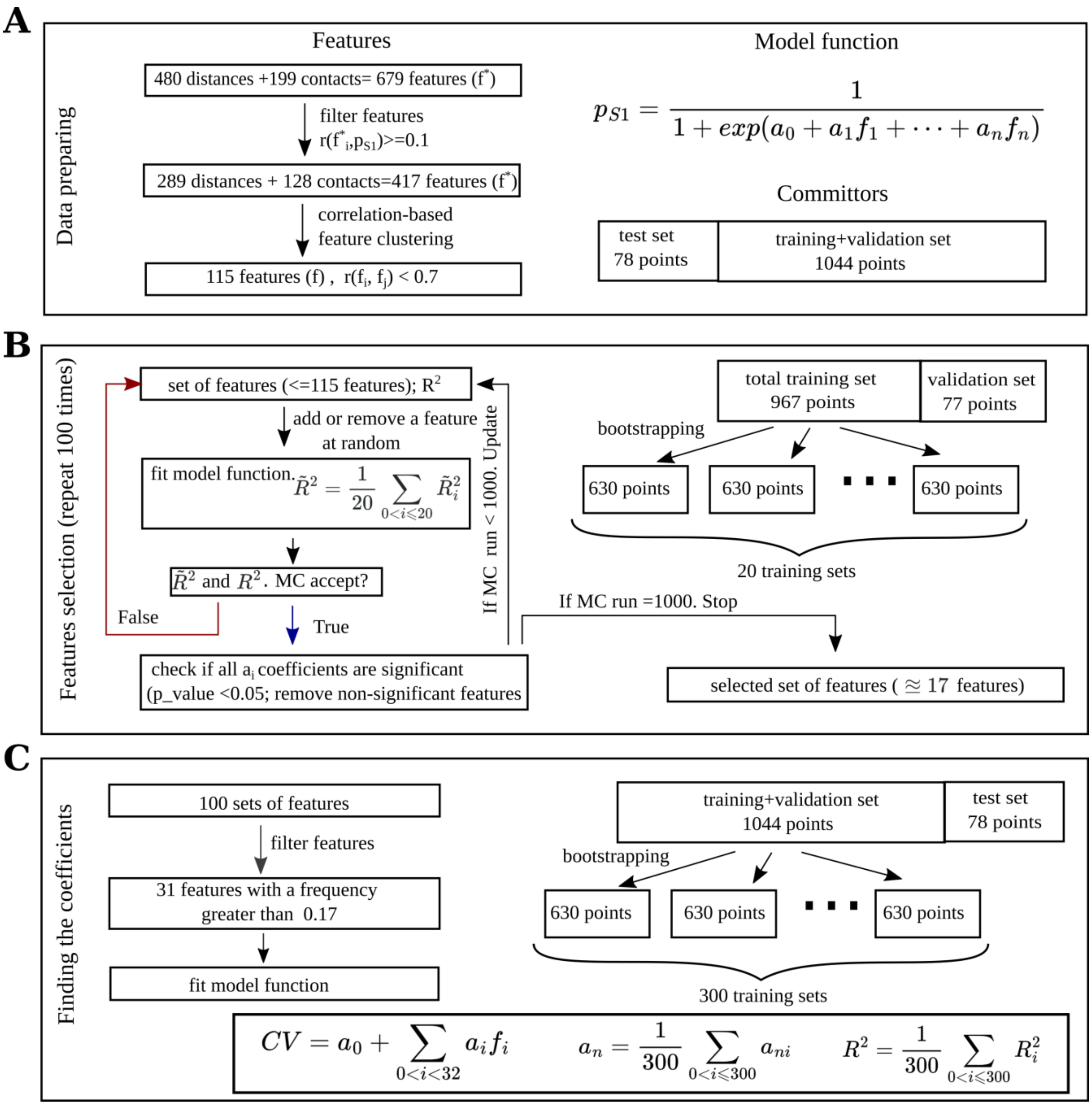


**Figure S3.** Methodology for the collective variable optimization. **A.** The initial set of features is subjected to elimination of multicollinearity. The total set of committor values is divided into a 78-point test set and a 1044-point remainder set. The form of the model logistic function is shown. **B.** The selection of features is conducted via a Monte Carlo procedure, with the objective of maximizing the adjusted $R^2$ value. The 1,044-point committor values set is divided into a validation set and a 967-point set. The latter is resampled 20 times using the bootstrap methodology, resulting in 20 training sets. For each combination of features, the model is fitted twenty times on different training sets, and the mean $R^2$ value is estimated on the validation set. This process is repeated 100 times. **C.** The 31 features with the highest frequency of occurrence were selected from the optimized combinations of features. The corresponding linear coefficients were then determined via fitting of the model function to committor values. This was achieved by fitting the model 300 times, with 300 training sets. Subsequently, the final adjusted $R^2$, MSE, and MAE values were calculated on a test set.

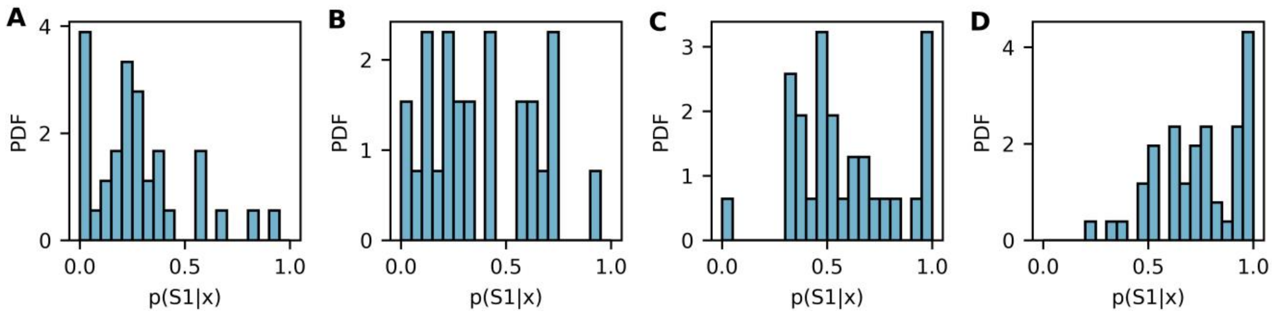


**Figure S4.** The histograms demonstrate the distributions of the committors for the various values of the optimized collective variable. The collective variable ranges from **A.** -1.25 to -0.75, **B.** -0.75 to -0.25, **C.** -0.25 to 0.25 (transition state), and **D.** 0.25 to 0.75.

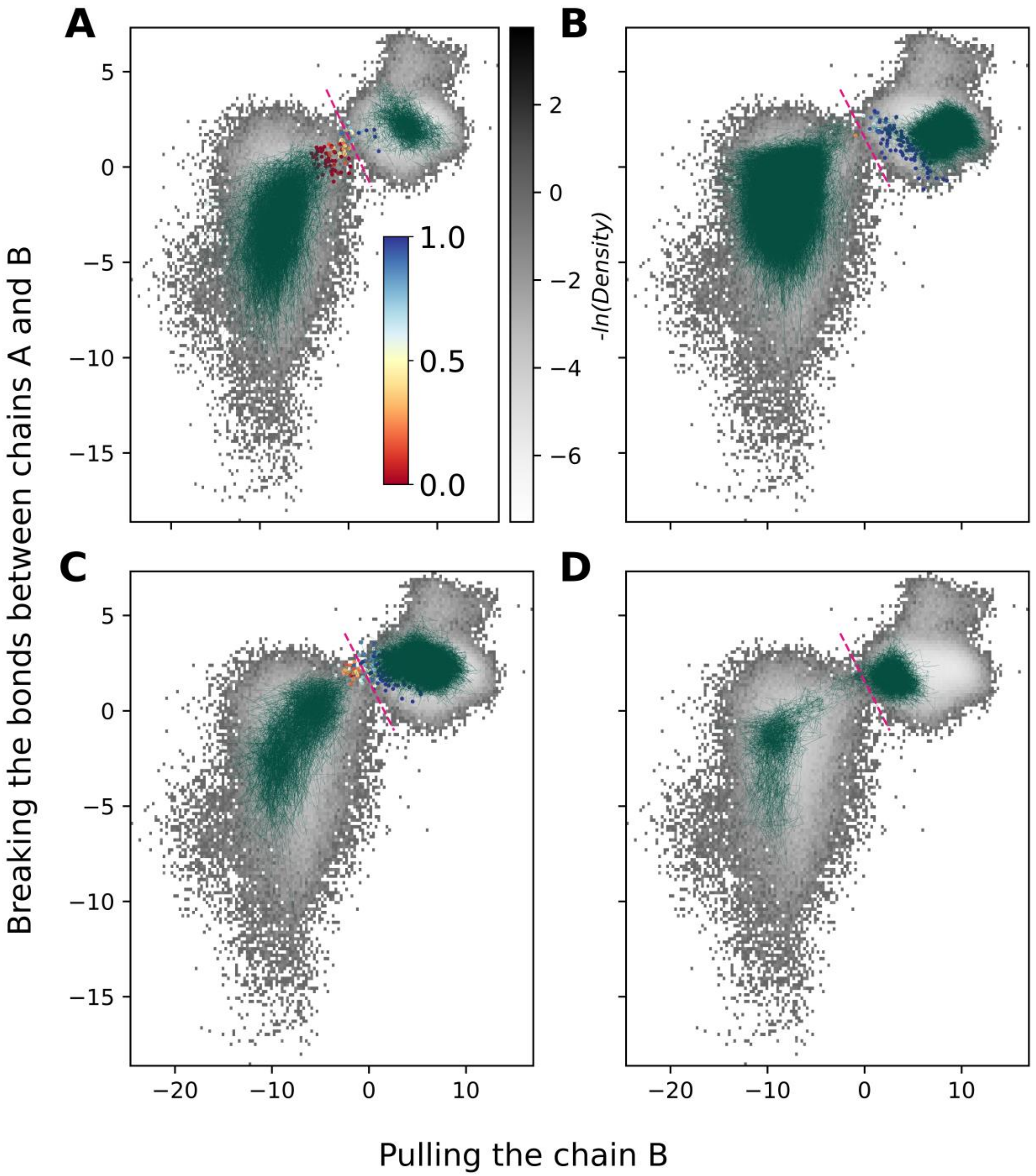


**Figure S5.** Individual trajectories plotted onto the space formed by the optimized collective variable divided into two parts. The gray landscape represents the density of all simulated paths, with the exception of replicas 4 and 11. All snapshots for which committors were computed are represented by dots colored according to the corresponding committor value. The pink dashed line indicates the optimal position of the transition state. The trajectories of each replica are displayed on the corresponding panels with a blue-green line: **A.** Replica 1, **B.** Replica 2, **C.** Replica 3, **D.** Replica 4.

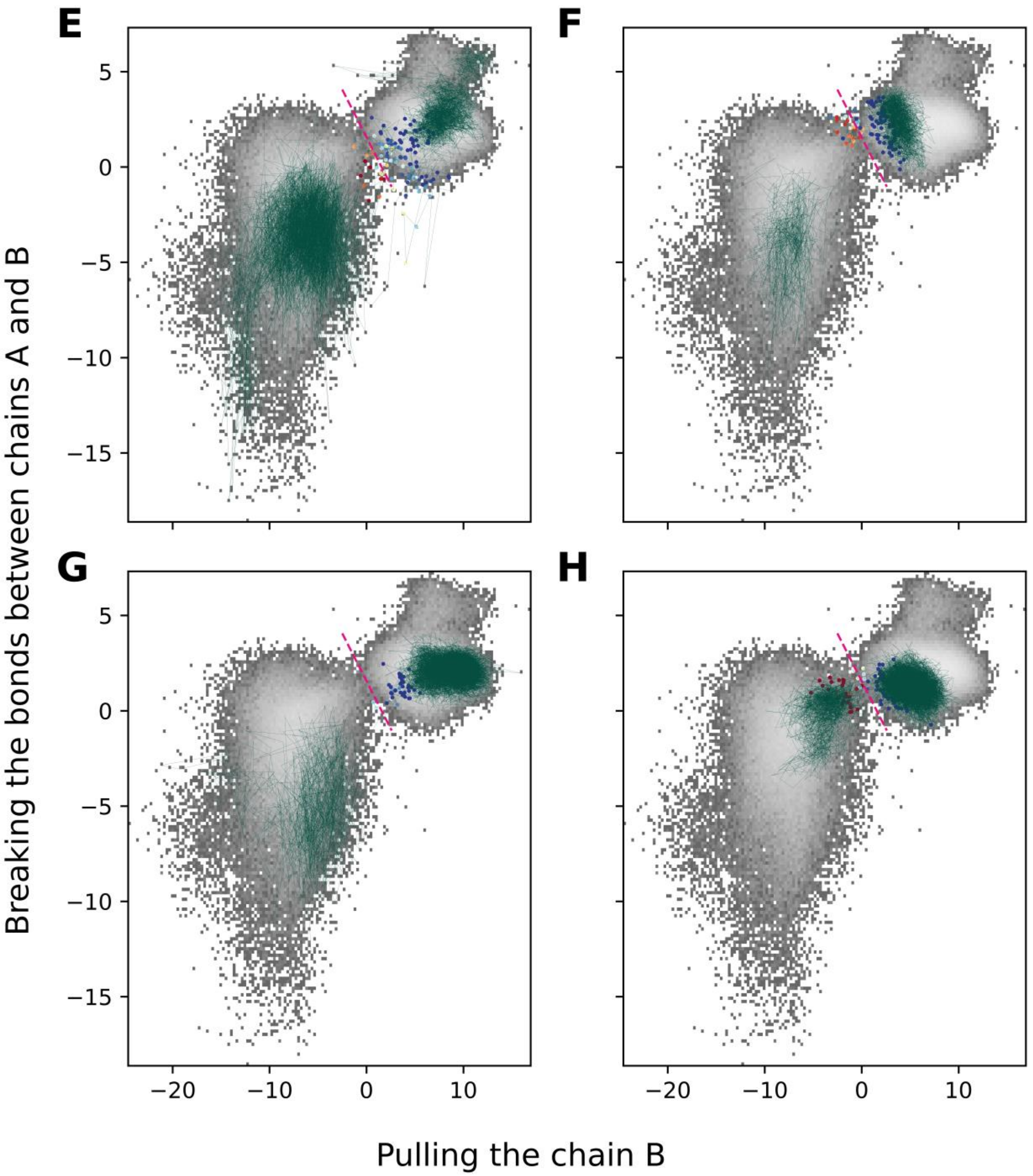


**Figure S5.** Continued. **E.** Replica 5, **F.** Replica 6, **G.** Replica 7a, **H.** Replica 7b.

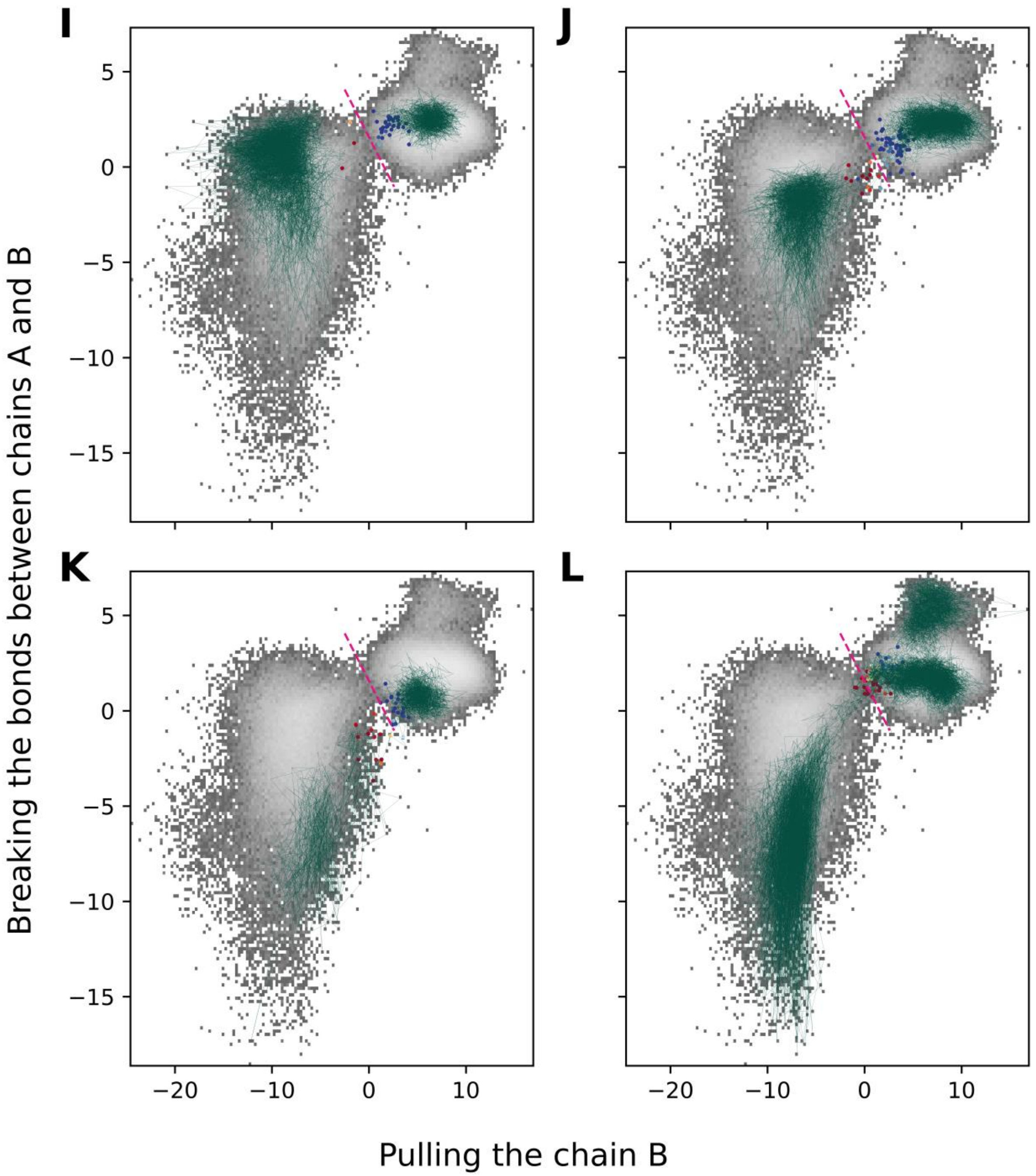


**Figure S5.** Continued. **I.** Replica 8 **J.** Replica 9, **K.** Replica 10, **L.** Replica 11.

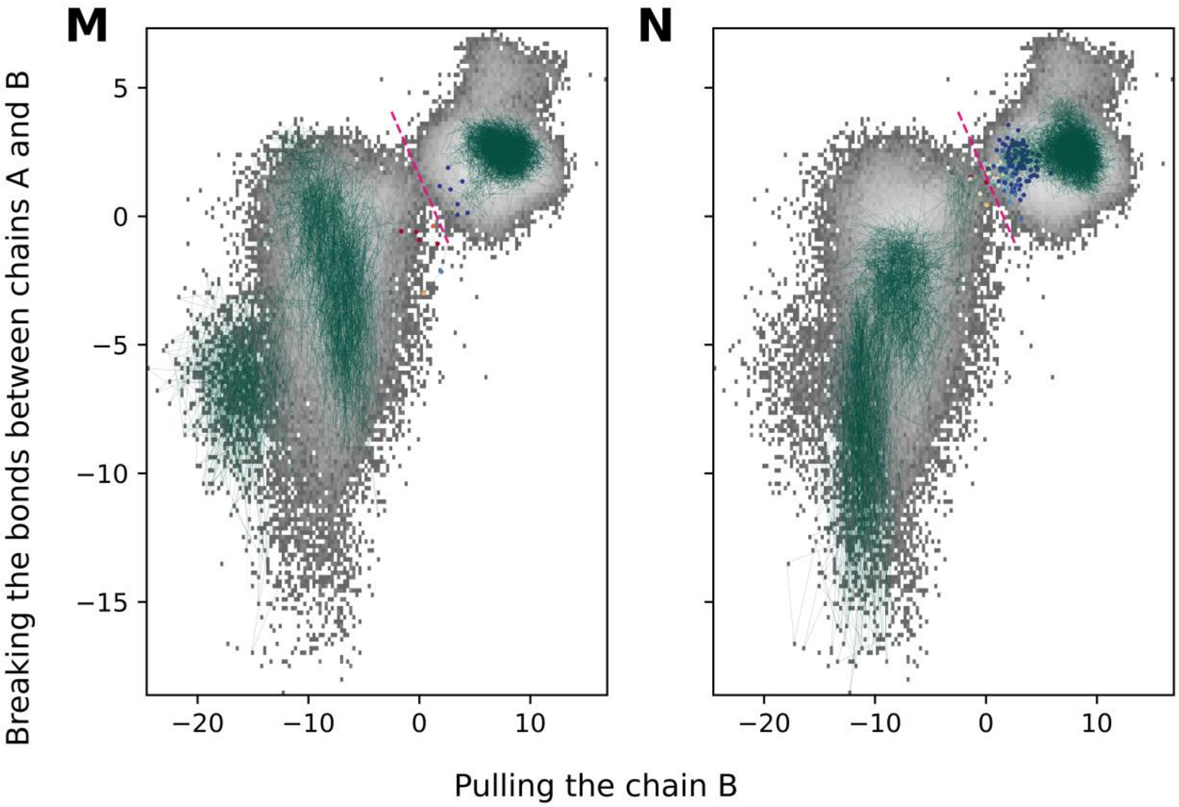


**Figure S5.** Continued. **M.** Replica 12a, **N.** Replica 12b.

S1. Contacts stabilizing the closed state of MscL

In this section, we will primarily describe the interactions between chain B and the other chains, since we have agreed that it is chain B that performs the conformational transition from the closed to the S1 state. It should be noted, however, that all of these interactions are present in all chains.

At zero tension, the closed state is stabilized by three discrete sites of contacts (Figure S6, S7a, Table S1), as well as interactions with lipid molecules in a lipid binding pocket formed by residues V22a, I82b and V86b, as proposed recently[1,2,3].

Table S1. Changes in the contact sites during the transition from the closed to the S1 state

| Sites of contacts | Location | Presence of the site in the closed state at zero tension | Presence of the site in the closed state at tension | Presence of the site in the transition state | Presence of the site in the S1 state |
|---|---|---|---|---|---|
| Site 1 (‘periplasmic’) | between the TM1 and TM2 helices of chain B | exists | almost disrupted | disrupted | disrupted |
| Site 2 (‘cytoplasmic’) | between the TM1 helices of chains A and B | exists | begins to weaken | partially disrupted | disrupted |
| Site 2 (‘cytoplasmic’) | between the TM1 helices of chains B and C | exists | exists | begins to weaken | partially disrupted |
| Site 3 | between the N-terminal helix of chain B and the TM2 helix of chain D | exists | exists | exists | exists; new contacts are formed between N-terminal helix of chain B and F80, A83, and F84 residues of chain D |
| Lipids in binding pocket | Acyl chains interacting with V22 residue of chain A and I82 and V86 of chain B | exists | begins to weaken | partially disrupted | disrupted |

We propose that the function of the ‘periplasmic’ and ‘cytoplasmic’ sites is to maintain the connection between adjacent channel ribs. The TM1 and TM2 helices of each chain are fastened at the top by means of the ‘periplasmic’ site, resulting in an almost orthogonal position of the ribs with respect to the membrane plane. Adjacent TM1 helices are bound near the cytoplasmic side of the membrane by the ‘cytoplasmic’ site. This site includes residues L17 and V21, which form a ‘vapor lock’ in the wild-type protein[4], suggesting that the 'cytoplasmic site' is primarily responsible for maintaining the closed state of the pore.

Considering the position of the sites, it can be assumed that the ‘periplasmic’ site resists forces applied to the periplasmic tension sensor (L72) and the "cytoplasmic" site resists forces applied to the "cytoplasmic" tension sensor (N-terminal helix).

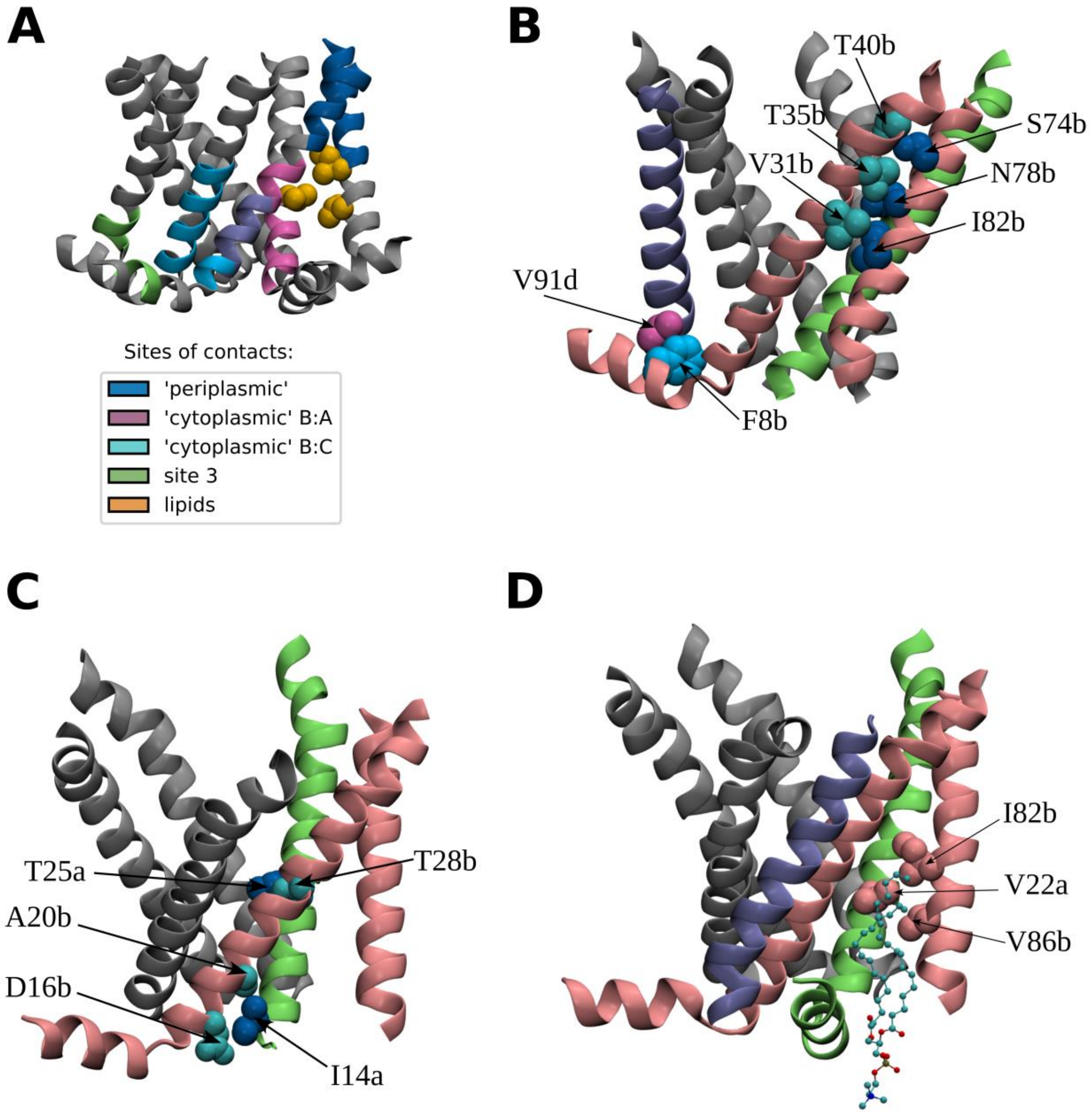


**Figure S6.** The closed state of MscL at zero tension. **A.** Relative position of all sites of contacts **B.** The 'periplasmic' contact site, represented by T40b:S74b, T35b:N78b, and V31b:I82b contacts, and the third site exemplified by the F8b:V91d contact. Chain B is highlighted in pink, the TM1 helix of chain A is shown in lime, the TM2 helix of chain D is highlighted in ice blue, and the remaining chains are shown in gray. **C.** The 'cytoplasmic' contact site between chains A and B, represented by T25a:T28b, I14a:A20b and I14a:D16b contacts. The chains are colored in the same pattern as in panel b, but the TM2 helix of chain D is omitted. **D.** Lipid binding pocket formed by the I82 and V86 residues from chain B and the V22 residue from chain A in the closed state at zero tension. The lipid molecule occupying the pocket is shown with a ball-and-stick model.

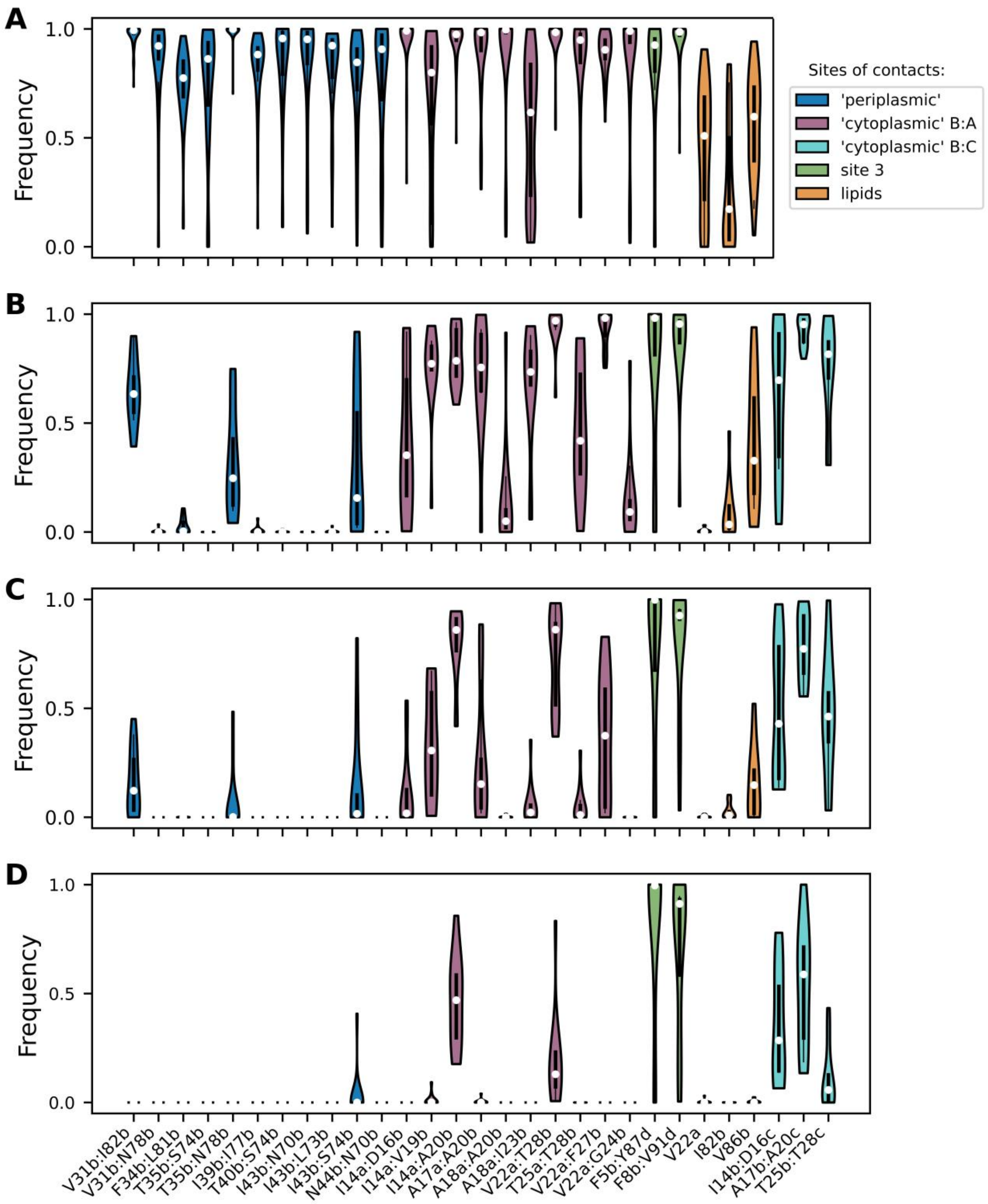


**Figure S7.** Frequencies of contacts stable in the closed state at zero tension during the transition of the chain B to the S1 state: **A.** closed state at zero tension, **B.** closed state under tension, **C.** transition state, defined as a range of the optimized collective variable from -5.4 to 2.4, **D.** S1 state. To plot the contact frequencies, two residues were considered to be in contact if the minimum distance between any two heavy atoms of the respective residues was less than 4.5 Å. The white circles represent the medians, the black thick lines extend from the first to the third quartile of the distributions.

Our simulations show that in the beginning of the transition applied tension has the most pronounced effect on the periplasmic side of the channel. Tensile forces not only effectively disrupts the ‘periplasmic’ site (fig. S7b), but also separates the two adjacent ribs, leading to the formation of the so-called 'funnel-shaped' structure[5]. We designated this structure as a closed state under tension, and as discussed in the main text, there is a free energy minimum corresponding to this state. Since, the majority of the ‘periplasmic’ site contacts are destroyed before reaching the transition state, their loosening does not correlate with the committor values and the disruption of these contacts does not contribute to the reaction coordinate connecting the closed and S1 states.

The conformational changes on the cytoplasmic side of the channel are relatively minor in the initial stage of the transition. However, the "cytoplasmic" site between chains A and B becomes weakened and further destroyed by the time the S1 state is reached (Fig. S7b-d). All the contacts belonging to this site contribute to the reaction coordinate. The 'cytoplasmic' site between chains B and C remains almost unchanged before reaching the reactive path region between the closed state under tension and the S1 state (Fig. S7b-d), but weakens significantly in the S1 state. The loosening of the contacts I14b:D16c, A17b:A20c, T25b:T28c has the highest correlation coefficient with the committor values.

The occupancy of the lipid binding pocket by the acyl chain begins to decrease in the closed state under tension. In the S1 state, there are no lipids in contacts with the pocket-forming residues (Fig. S7b-d). The delipidation of the lipid binding pocket contributes to the reaction coordinate and is described in detail in section S3.2.

S2. The transition of MscL from a closed to an S1 state is dependent upon the presence of lipids.

Attempts to distinguish between the closed and S1 states of MscL revealed that this was not feasible in all replicas when using solely the coordinates of protein atoms. An investigation into potential factors that could enhance the distinction between the closed and S1 states identified stable interactions between lipids and three MscL residues, specifically V22, I82, and V86, forming a kind of binding pocket for fragments of lipid acyl chains. These interactions are sufficiently frequent in the closed state under zero tension and still persist in the closed state when tension is applied. However, these interactions never occur in the S1 state. It seems that the elimination of lipids from a protein surface is a stochastic process that can be made more probable by the applied tension. If the lipids remain in the binding pocket formed by the V22 residue of chain A and the I82 and V86 residues of chain B for a longer period of time, they may play a crucial role in preventing the transition of MscL into the S1 state. The most illustrative example of this phenomenon is the Replica 7b, which is elucidated in Figure S8a. In the figure, two collective variables are introduced, both of which are the first principle component calculated on a set of contacts. However, the initial set comprises solely protein residues, whereas the subsequent set incorporates additionally the contacts between lipids and the V22a:I82b:V86b binding pocket. The two histograms illustrate the density of states along these variables. The peak on the right corresponds to the closed state at tension, while the peak on the left corresponds to the S1 state. It can be clearly seen that the peaks are separated only when interactions with lipids are included. The molecular mechanism behind the phenomenon is as follows (Fig. S8b). As discussed in the main text, during the transition to the S1 state, helix TM1 of chain B is subjected to a pulling force, resulting in its displacement in relation to the ‘rib’ formed by helix TM1 of chain A and helix TM2 of chain B. As a consequence of this displacement, the I39 residue from the TM1 helix of chain B moves in closer proximity to residues A85, V86, L89, and V90 from the TM2 helix of chain B, thereby forming stable contacts with them. However, the lipid acyl moiety occupying the V22a:I82b:V86b binding pocket precludes the displacement of the TM1 helix, as there is no longer available space to accommodate I39. In Replica 7b, this factor constituted the sole impediment to the transition to state S1 for a certain period of time.

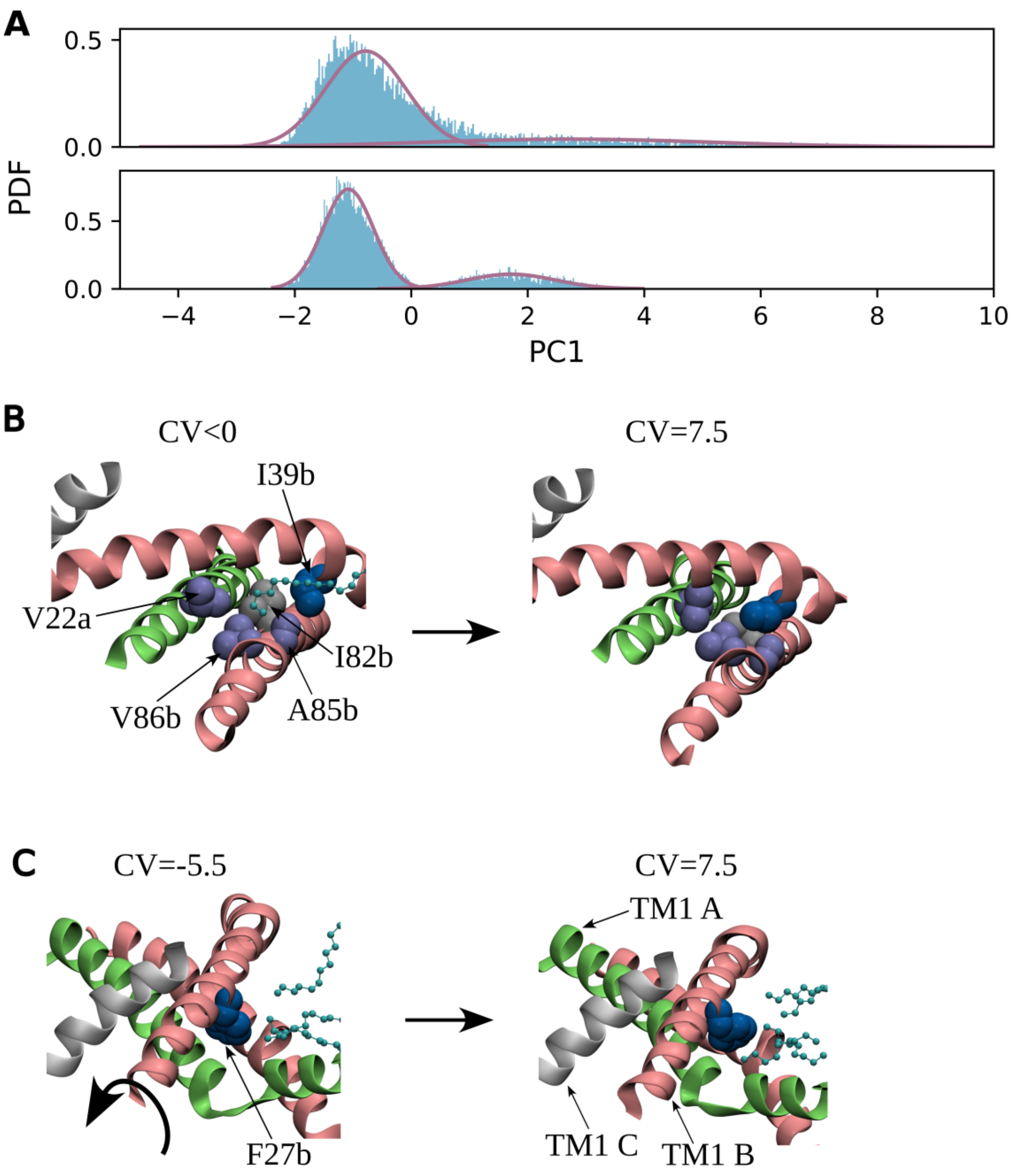


**Figure S8.** Impact of lipids on the transition of MscL to the S1 state. **A.** The probability density functions of two collective variables for Replica 7b. Both variables represent a first principle component calculated on a set of contacts. However, the top set comprises solely protein residues, whereas the bottom set incorporates additionally the contacts between lipids and the V22a:I82b:V86b binding pocket. The peak on the right corresponds to the closed state at tension, while the peak on the left corresponds to the S1 state. **B.** The left-hand representation illustrates the positioning of the lipid acyl chain within the V22a:I82b:V86b binding pocket in the closed state, with the applied tension. The right-hand representation depicts the I39 residue of the TM1 helix of chain B, which occupies the same pocket and forms contacts with the A85 and V86 residues in the S1 state. **C.** Upon reaching the S1 state, the F27 residue of chain B increases contacts with lipid acyl moieties, which is accompanied by a slight clockwise rotation of the TM1 helix.

The analysis of correlations between molecular features and committor values revealed the existence of an additional mechanism involving lipids in the control of MscL activation. In the S1 state, the F27 residue of chain B was observed to increase its contacts with lipids, accompanied by a slight clockwise rotation of the TM1 helix of chain B around its axis (Fig. S8c). While a comprehensive analysis of the causative and consequential relationships remains to be performed, it may be hypothesized that the favorable contacts with lipids cause the helix rotation, rather than being a consequence of it. The clockwise rotation of the TM1 helix was previously reported in the literature and is presumed to be a requisite step in the MscL gating[6].

S3. Conductance of the MscL states

We quantified the conductance of MscL states at -100 mV membrane potential and 1 M KCL concentration. The physiological concentration of salt is approximately 10 times lower, which would imply that the "true" conductance should be 10 times lower as well.

The closed state under tension is leaky with a conductance of 0.3 ± 0.1 nS. We hypothesized that leakage occurs exclusively in the L17A V21A mutant. At the very least, our simulations of the closed state of the wild-type MscL under tension did not indicate the presence of any conductance.

Conductance of the S1 state is $2.7 \pm 0.4$ nS . This value is approximately ten times lower than the open state conductance (2.5-4 nS)[7,8]. This allows the assumption to be made that the pore area of the S1 state is approximately ten times lower than that of the open state pore. It is noteworthy that the conductance of MscL does not appear to correlate with committor values. While a comprehensive analysis has yet to be performed, one potential explanation is that in the region of the barrier, alterations in the pore area are relatively minor, and the primary movement of the TM1 helix of chain B, which results in an increase in the pore size, occurs subsequent to the barrier having been crossed.

SM REFERENCES

1. Kapsalis, C; Wang, B; Mkami, H. El.; Pitt, S. J.; Schnell, J. R.; Smith, T. K; Lippiat, J. D.; Bode, B. E.; Pliotas, C. Allosteric activation of an ion channel triggered by modification of mechanosensitive nano-pockets. *Nat. Commun.* 2019, 10 (1), 4619. DOI: 10.1038/s41467-019-12591-x.

2. Kapsalis. C; Ma, Y; Bode, B. E.; Pliotas, C. In-Lipid Structure of Pressure-Sensitive Domains Hints Mechanosensitive Channel Functional Diversity. *Biophys J.*, 2020, 119 (2), 448-459. DOI: 10.1016/j.bpj.2020.06.012

3. Wang, B.; Lane, B. J.; Kapsalis, C.; Ault, J. R.; Sobott, F.; Mkami, H. El., Calabrese, A. N.; Kalli, A. C.; Pliotas, C. Pocket delipidation induced by membrane tension or modification leads to a structurally analogous mechanosensitive channel state. *Structure*, 2022, *30* (4), 608–622.e5. DOI: 10.1016/j.str.2021.12.004

4. Anishkin, A.; Akitake, B.; Kamaraju, K.; Chiang, C. S.; Sukharev, S. Hydration properties of mechanosensitive channel pores define the energetics of gating. *J. Phys.:Condens. Matter*, 2010, 22 (45), 454120. DOI: 10.1088/0953-8984/22/45/454120

5. Betanzos, M.; Chiang, C. S.; Guy, H. R.; Sukharev, S. A large iris-like expansion of a mechanosensitive channel protein induced by membrane tension. *Nat. Struct. Biol.*, 2002, 9 (9), 704-710. DOI: 10.1038/nsb828.

6. Iscla, I.; Levin, G.; Wray, R.; Reynolds, R.; Blount, P. Defining the physical gate of a mechanosensitive channel, MscL, by engineering metal-binding sites. *Biophys J.*, 2004, *87* (5), 3172–3180. DOI: 10.1529/biophysj.104.049833

7. Yang, L. M; Zhong, D.; Blount, P. Chimeras reveal a single lipid-interface residue that controls MscL channel kinetics as well as mechanosensitivity. *Cell Rep.,* 2013, 3 (2), 520-527. DOI: 10.1016/j.celrep.2013.01.018.

8. Moe, P. C.; Blount, P.; Kung, C. Functional and structural conservation in the mechanosensitive channel MscL implicates elements crucial for mechanosensation. *Mol. Microbiol.,* 1998, 28 (3), 583-592. DOI: 10.1046/j.1365-2958.1998.00821.x.
